# The potential of mycelium from mushroom-producing fungi in alternative protein production: a focus on fungal growth, metabolism, and nutrition

**DOI:** 10.1101/2025.08.29.673026

**Authors:** Jasper Zwinkels, Stef van Oorschot, Oscar van Mastrigt, Eddy J. Smid

## Abstract

The growing need for high-quality protein with minimal environmental impact necessitates the expansion of alternative proteins on the market. One area with great opportunity for expansion lies in the phylogenetic diversity of the fungal kingdom. Diversifying the use of fungal species, by assessing species from the phylum of mushroom-producing fungi (Basidiomycota) in solid-state fermentation, could open new avenues to foods with improved nutritional and sensorial properties. To assess these properties, we first determined the potential of basidiomycetes to ferment and colonize cereals and legumes. A phylogenetically diverse selection of eight species of basidiomycetes was analyzed on their radial growth speed and biomass yield. The best performing species were successfully fermented on brown rice (high starch), brewer’s spent grain (high fiber, high protein), and lupin (high protein, high fiber and high fat), and compared to *Rhizopus microsporus* var. *oligosporus*. Large variation in performance was observed between the different basidiomycetes on the three substrates in terms of biomass formation and metabolic behavior. The presence of an easily accessible carbon source, such as starch was needed to prevent deamination and thereby loss of valuable protein. With the correct formulation, basidiomycetes could fully ferment and colonize the substrate, thereby increasing the overall protein content and degrading the anti-nutritional factor phytic acid up to 80%. These results provide a methodology for screening of fungal species and substrates and demonstrate that basidiomycetous mycelia represent a promising source of phylogenetic diversity for novel food fermentations.

**Highlights:**

‐ Brown rice, brewer’s spent grain, and lupin were fermented with basidiomycetes.
‐ Readily available carbon prevents protein catabolism during fermentation.
‐ Fungal growth was analysed visually, thermally and through metabolic indicators.
‐ Basidiomycetes increased the protein content and reduced phytate more than Rhizopus.

## 1 Introduction

Animal-based foods account for 53% of food-related anthropogenic greenhouse gas emissions and its production occupies 40% of the habitable land (Poore & Nemecek, 2018; Xu et al., 2021). Adoption of more non-animal proteins in our diets can greatly reduce the environmental impact of food production (Jarmul et al., 2020). This growing need for sustainable food production has intensified the search for alternative proteins with minimal environmental impact while meeting global nutritional demands.

A promising approach to more sustainable food production is fermentation using filamentous fungi, an ancient food processing technique responsible for products like tempeh and koji (Strong et al., 2022). Despite its long history, fungal fermentation-based foods have yet to gain widespread acceptance, largely due to factors such as food conservatism, costs, nutritional quality, and taste (Tso et al., 2020). Current research primarily focuses on optimizing fermentation efficiency and improving existing production systems (Ahmad et al., 2022). However, commercially available fungal-fermented foods are derived from a narrow range of fungal species—primarily from six genera across two fungal phyla (Table 1). While altering substrate composition and growth conditions allows for some product diversification (Wang et al., 2023), expanding the diversity of fungi used in fermentation could unlock a wide range of novel food applications, with potentially improved nutritional and sensorial qualities.

**Table 1.** Commonly available food products produced with filamentous fungi.

| Phylum | Genus | Type of fermentation | Food products |  |
| --- | --- | --- | --- | --- |
| Ascomycota | *Aspergillus* | Solid-state | Koji, sake, soy sauce, miso | (Allwood et al., 2021; Zwinkels et al., 2023) |
| Mucoromycota | *Rhizopus* | Solid-state | Tempeh | (Zwinkels et al., 2023) |
| Ascomycota | *Fusarium* | Submerged | Mycoprotein | (Finnigan, 2011) |
| Ascomycota | *Penicillium* | Solid-state | Mold-ripened cheese | (Yin et al., 2020) |
| Ascomycota | *Neurospora* | Solid-state | Oncom | (Starzynska-Janiszewska et al., 2017) |
| Mucoromycota | *Actinomucor* | Solid-state | Sufu | (Han et al., 2004) |

A particularly promising, yet underexplored group is the phylum Basidiomycota, which comprises approximately 31,000 species, many of which produce edible fruiting bodies (mushrooms). These species are promising, firstly, because their fruiting bodies are already part of human diets and are valued for their umami-rich flavor and meat-like texture (Mau, 2005; Yuan et al., 2022). This could make the consumption of basidiomycetous mycelium more likely to be safe and tasteful than other unleveraged fungal species. Secondly, the mycelium of basidiomycetes generally has a higher nutritional value (e.g. more protein and more bioactive compounds), and mycelium production is faster (days instead of months), and more efficient than producing fruiting bodies (Muswati et al., 2021; Papaspyridi et al., 2010). Thirdly, basidiomycetes can grow on agricultural side streams, particularly lignocellulosic materials, offering a sustainable approach to food production (Ahlborn et al., 2019). Lastly, many species of basidiomycetes greatly reduce anti-nutritional factors, such as phytic acid, and to a larger extent than fungi conventionally used in fermentation, such as Rhizopus sp. (da Luz et al., 2013; Vong et al., 2018). For these reasons, they are strong candidates for alternative protein production and leveraging the genetic diversity found in Basidiomycota could open new avenues for the development of fungal-based protein products with improved flavor and texture and with nutritional and environmental benefits.

Despite their potential for food applications, research on Basidiomycota has primarily focused on growth on lignocellulosic side streams unfit for human consumption, which yields mycelial products intended for animal feed rather than human food. There are no studies investigating the potential of basidiomycetous mycelium grown on food-grade substrates as novel fermented food. This represents a significant knowledge gap in the field, as leveraging fermentation driven by basidiomycetes could open new possibilities for alternative protein production with improved sensory and nutritional properties.

To establish the nutritional and sensory benefits of solid-state fermentations (SSF) with members of the phylum of Basidiomycetes, their growth and metabolism on food-grade substrates should first be assessed because food-grade substrates differ in composition from lignocellulosic materials.

To assess the ability of basidiomycetes to metabolize and grow on food-grade substrates, we selected substrates with a diverse nutritional composition, namely brown rice, brewer’s spent grain (BSG) and lupin. Specifically, brown rice is high in starch and low in protein, BSG is high in fiber and high in protein, and lupin is high in fiber, protein, and fat (table 2). These substrates serve as models for the diverse nutritional composition of fungal fermentation substrates.

**Table 2.** Chemical composition of substrates used in this study.

| Component<br>(g/100 g DW) | Brown rice<br>(USDA, 2019) | Brewer’s spent grain<br>(Castro & Colpini, 2021; Jin et al., 2022) | White lupin<br>(Martínez-Villaluenga et al., 2006; Večerek et al., 2008) |
| --- | --- | --- | --- |
| Water<sup>*</sup> | 63.8<sup>1</sup> | 78.6<sup>1</sup> | 65.6<sup>1</sup> |
| Carbohydrates | 86.2 | N.D. | 30.3 |
| Starch | 83.5 | 1-12<sup>2</sup> | 5.9 |
| Crude fiber | 5.4 | 29.9 | 23.4 |
| Protein | 8.6<sup>1</sup> | 17.4<sup>1</sup> | 36.1<sup>1</sup> |
| Fat | 3.3 | 1.6 | 15.0<sup>1</sup> |

<sup>1</sup>Results obtained in this study.
<sup>2</sup>Varies based on brewing efficiency.
<sup>\*</sup>(g/100 g)
N.D. = No Data

In addition to a broad selection of substrates, a taxonomically diverse selection of species of the phylum of Basidiomycota is important to cover a broad range of metabolic growth strategies. To build upon existing knowledge and produce a product safe for human consumption, species that are commercially cultivated and produce edible mushrooms were selected. While these fungi are well-characterized in terms of cultivation for mushroom production, their growth dynamics on food-grade substrates remains unclear, necessitating robust biomass estimation methods.

The ability of filamentous fungi to colonize a substrate has been notoriously difficult to monitor, due to the intertwined nature of the mycelium and the substrate (Steudler & Bley, 2015b). Therefore, many indirect methods, such as visual growth observation, heat production measurements, and biomarker estimation, such as ergosterol and glucosamine, have been employed to estimate fungal biomass (Steudler & Bley, 2015a). A combination of methods could further improve the precision of fungal biomass estimations. Furthermore, understanding the metabolic strategies fungi employ to colonise different substrates is valuable, as this knowledge can inform the systematic selection of fungal species for substrates with compatible chemical compositions.

In this study, we aim to evaluate the ability of commercially cultivated species of the phylum of Basidiomycota to colonize and metabolize food-grade substrates with distinct nutritional properties. Specifically, we assessed (i) mycelial growth using a combination of biomass estimation methods, (ii) fungal metabolism through changes in nutrient composition post-fermentation, and (iii) potential nutritional benefits, including the degradation of anti-nutritional factors. By addressing these three aspects, this research provides insights into the feasibility of a novel solid-state fermentation using mycelium of basidiomycetes as an alternative protein source

## 2 Materials & Methods

### 2.1 Substrates

Brown Basmati rice (*Oryza sativa*) from Molen de Vlijt (Wageningen, the Netherlands), white halved lupin seeds (*Lupinus albus*) from France (Inveja Food) and Brewer’s spent grain (*Hordeum vulgare*) from Stadsbrouwerij Veenendaal (Veenendaal, the Netherlands) were used as substrates in solid-state fermentation. Brewer’s spent grain (BSG) consisted of malted pilsner barley.

### 2.2 Fungal species

*Lentinula edodes* CBS 134.85 (Shiitake), *Flammulina velutipes* CBS 347.74 (wild enoki), *Stropharia rugosoannulata* CBS 288.85 (garden giant), *Coprinus comatus* CBS 150.39 (Shaggy mane), *Schizophyllum commune* CBS 462.62 (Splitt gill fungus), *Volvariella volvacea* CBS 312.84 (Straw mushroom) and *Pycnoporus cinnabarinus* CBS 311.33 and *Rhizopus microsporus* var. *oligosporus* CBS 338.62 were obtained from the public culture collection of Westerdijk Fungal Biodiversity Institute (Utrecht, The Netherlands). *Pleurotus pulmonarius* (M2345) (sold as *Pleurotus sajor-caju*) was obtained as mother culture on a petri dish from Mycelia BVBA (Deinze, Belgium).

### 2.3 Inoculum preparation

#### 2.3.1 Basidiomycete inoculum preparation

A square (1×1 cm) agar cube with mycelium was transferred from slant or plate to the middle of a new malt extract agar plate (MEA, Oxoid) and incubated at their optimal temperature until the plate was (almost) fully covered. A square (1×1 cm) agar cube was then transferred to a 500 mL Erlenmeyer containing 200 mL sterile malt extract broth (MEB, Oxoid) and subsequently incubated in shaking incubators at their optimal temperature at 180 rpm. Optimal temperatures determined by assessing the radial growth, described in section 2.5. When fully grown, the mycelium was broken apart using a sterile 50 mL pipet and 40 mL was transferred aseptically to a 50 mL Greiner tube. The suspension was washed 3 times using physiological saline solution (0.9% NaCl) and concentrated 2 times to obtain 20 mL mycelium suspension.

#### 2.3.2 Rhizopus inoculum preparation

*Rhizopus microsporus* inoculum was prepared as fungal spore suspension according to the method described by Wolkers – Rooijackers et al. (2018).

### 2.4 Fungal growth rate on agar plate

The pH of malt extract agar (MEA, Oxoid) plates was altered by addition of 1 M HCl (Merck, Burlington, MA, USA) or 1 M NaOH (Merck) before autoclaving. Radial growth of fungi was estimated by aseptically transferring a 1×1 cm cube of agar from a fully grown agar plate to the center of a fresh plate of MEA. Plates were incubated at 25, 27, 30 and 33°C and at the optimal temperature at pH 3.5, 4, 5, 6 and 7 for up to 8 days. Radial growth (cm) was measured daily as the diameter of the colony minus the initial size of the inoculation cube.

### 2.5 Yield & dry weight loss estimation in liquid culture

Maximum yield and dry weight reduction were estimated in a liquid culture of malt extract broth (MEB, Oxoid) acidified to the optimal pH. MEB was sterilized and the dry matter content was measured using a moisture analyzer MA100 (Sartorius, Germany) at 105°C until a constant weight was reached. A small piece of mycelium was scraped off a fully grown agar plate and broken apart before aseptically adding it to a dry, pre-weighed, and sterile Erlenmeyer (150 mL) containing 50 mL MEB. Erlenmeyers were incubated at optimal temperature, determined by radial growth on agar plate, described in section 2.4, at 180 RPM agitation. Incubation occurred until stationary phase was reached, which took between 12 and 19 days, depending on the species. After incubation, the Erlenmeyers were weighed, after which mycelium was filtered through a pre-weighed Whatman filter paper (cellulose, 190 µm x 125 mm, pore size 8 µm). Dry matter content of the filtrate was determined in the moisture analyzer. Thereafter, the mycelium was washed with around 1 mL demineralized water and dried to a constant weight at 80°C. The weight loss due to evaporation of water in the filter paper was accounted for by determining the weight loss of an empty filter paper. This filter paper was weighed, then rinsed with demi water and hence dried at 80°C. Filter papers were weight on an analytical balance. The maximum yield and dry weight loss were calculated according to equations 1, 2, 3 and 4.

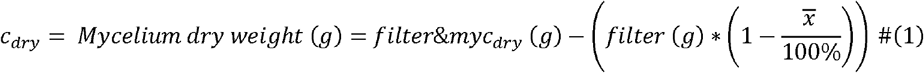

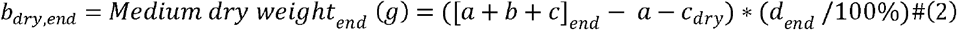

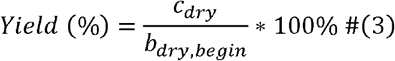

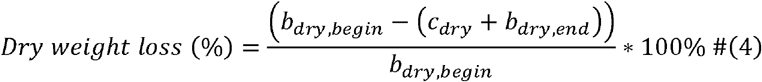

*a = Erlenmeyer (g)

*b = Medium (g)

*c = Mycelium (g)

*d = Medium dry matter content (%)

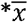 = average filter weight loss during drying (%) (n=5)

### 2.6 Solid-state fermentation

Substrates (500 g) were soaked for 16 hours at room temperature (20°C) in plastic jars with substrate to tap water (w/w) ratio of 1:3. The substrate water mixture was acidified to pH 4.0 using 1 M lactic acid. After soaking the water was drained and substrates were rinsed using tap water for 30 s. Substrates were divided over filter boxes (80 g brown rice and BSG, or 100 g lupin, per box; O118/50, Microbox polypropylene fermentation container, Sac O_2_, Belgium) and closed with filter lids (OD118, red filter, Sac O_2_, Belgium). Water was added for rice, and lupin (1:1 and 0.4:1 (w/w)), while BSG pressed to remove excess water. Filter boxes were packaged in waste bags and autoclaved for 15 min at 121°C. After sterilization, condensation was removed by air drying the boxes under laminar flow. Substrates were inoculated by addition of 5 mL of mycelium suspension (0.1 g dry mycelium) per 100 g substrate or 1 mL of spore suspension (10^4^ viable spores/gram) per 100 g substrate, respectively. Samples were incubated at their optimal growth temperature (determine by radial growth rate, described in section 2.4) for 8 and 2 days for fermentations with basidiomycetes or Rhizopus, respectively. Uninoculated autoclaved samples were used as control (Unfermented). All fermentations were conducted in triplicate.

### 2.7 Sample preparation

After fermentation, the fermented cake was cut into 8 identical slices and pieces from all sides of the sample were taken, frozen with liquid nitrogen and ground to a fine powder using a spice grinder (Krups F203, Solingen, Germany). Homogenized samples were immediately stored in a urine container (VWR international, ML15) at -20°C until further analysis.

### 2.8 Fungal biomass estimation in solid-state

#### 2.8.1 Visual appearance

Visual appearance was assessed by taking pictures of the solid-state fermented samples directly after fermentation.

#### 2.8.2 Ergosterol quantification

Fungal biomass was estimated through ergosterol in the SSF sample and calibrated based on ergosterol content in dry mycelium cultivated on agar plates. Ergosterol was extracted from the SSF samples as described by Sae-Tun et al. (2020), which used a variation on the method of Gong et al. (2001). Ergosterol concentrations were translated to fungal biomass as described by Zwinkels et al. (2023), with the adaptation that for basidiomycetes a mycelium suspension of 1 mL was used instead of a 100 µL fungal spore suspension.

#### 2.8.3 Glucosamine and amino acid content

Fungal biomass was calculated through glucosamine based on glucosamine concentration in fermented samples in relation to that in pure mycelium cultivated in broth. Mycelium was produced according to the procedure described in section 2.5. Unfermented and fermented samples, and mycelium were frozen using liquid nitrogen and grounded to a fine powder using a coffee grinder (Krups F203) before being lyophilized.

Before amino acid (AA) and glucosamine determination procedures, unfermented and fermented lupin samples were defatted (Thiex, 2009). The total amino acid content and glucosamine content were determined according to ISO13904 (ISO, 2005b) and ISO13903 (ISO, 2005a) procedures. In brief, cysteine and methionine were extracted by oxidization before hydrolysis of the protein. For all other AAs and glucosamine, no oxidization step was required. The AAs and glucosamine were separated by ion exchange chromatography and determined by reaction with ninhydrin, using photometric detection at 570 nm and 440 nm for proline. Tryptophan was determined by reversed-phase C18 HPLC with fluorescence detection. These chemical analyses were executed in duplicate and when the coefficient of variation was >5% analyses were repeated.

### 2.9 Fungal metabolism

#### 2.9.1 Temperature profile

The temperature profile during SSF was measured using a temperature logging iButton (iButtonLink, Whitewater, WI, USA), which was placed in the core of the sample. Temperature was measured at 10-minute intervals for the duration of the fermentation period. Heat production of fungal metabolism was derived from temperature data and expressed as °C above incubator temperature.

#### 2.9.2 pH

The pH of samples was measured in unfermented and fermented samples according to the following procedure. Ten grams of homogenized sample was weighed into a 50 mL Greiner tube to which 20 mL demineralized water was added. Tubes were vortexed for 20 seconds before pH was measured using a pH meter (MeterLab PHM240, Denmark).

#### 2.9.3 Dry matter loss

Dry weight loss during SSF was determined by subtracting the total dry weight of a fermented sample from the dry weight of the unfermented substrates plus the dry weight of the inoculum. Weights were recorded on an analytical balance and dry matter contents were determined in homogenized samples using a moisture analyzer MA100 (Sartorius, Germany).

#### 2.9.4 Sugars and anti-nutritional factors

Maltose and glucose and the anti-nutritional factors phytic acid and oxalic acid were analyzed using the same method as Scott et al. (2021), with some adaptations explained in brief. 0.1 gram of homogenized sample was weighed in a 15 mL greiner tube, to which 1 mL HCl (0.8 M) was added. Tubes were immediately put on ice, vortexed for 10 seconds and heated in a boiling water bath (100°C) for 1 min. Hereafter, the tubes were incubated at 30°C for two hours at 300 rpm. The tubes were cooled to room temperature and centrifuged at 14,000 g for 10 min (Pico 21 microcentrifuge, Thermo Scientific; Waltham, MA, USA). Samples were deproteinated by Carrez treatment and compounds were quantified by high-performance liquid chromatography (HPLC) as described by Scott et al. (2021) using an UltiMate3000 (Thermo Scientific; Waltham, MA, USA) equipped with an Aminex HPX-87H column (Biorad) kept at 40°C. The mobile phase was set at a flow rate of 0.6 ml/min and composed of 0.026 N H_2_SO_4_, with a runtime of 25 min. Sugars were detected by refractive index (RefractoMax520) and acids were detected by UV measurements at 220 nm. The calibration curve consisted of 0-20 mM maltose, 0-40 mM glucose, 0-80 mM oxalic acid and 0-4.0 mM phytic acid.

#### 2.9.5 Crude protein content

Nitrogen content was determined by DUMAS method with FlashSmart™ N/PROTEIN Element analyzer (Thermo Scientific) in accordance with AOAC, 2006.03 (2006). Crude protein contents were calculated using food-specific protein-nitrogen conversion factors proposed by Mariotti et al. (2008). Conversion factors were 5.45, 5.345 and 5.34 for brewer’s spent grain, lupin, and brown rice, respectively.

#### 2.9.6 Amino acid content

Procedures used in the determination of the total amino acid content are described under section 2.8.3. Aromatic amino acids (phenylalanine and tyrosine) could not accurately be determined using these hydrolysis methods and were therefore excluded from the total amino acid content.

#### 2.9.7 Ammonia content

The ammonia content was determined in a 5-fold diluted, homogenized sample, based on the method described by Scott and co-workers (2021).

### 2.10 Data analysis

Data generated by HPLC were analyzed in Chromeleon 7.3.1 (Thermo Scientific, MA, USA). Data was analyzed and figures were produced using Rstudio software v. 2024.12.0 (RStudio®, Boston, MA, USA). Significant differences indicated with different letters were determined using One-Way ANOVA with Tukey post hoc test. Significance level (p-value) was set at 0.05. All values presented are means of biological triplicates, unless stated otherwise.

## 3 Results

### 3.1 Growth dynamics of pure mycelium

To determine the growth speed of the selected basidiomycetes, mycelium was cultivated on malt extract agar (MEA) plates by placing a small square agar cube of fully grown agar in the center. Growth conditions were optimized for all fungal species by differentiating temperature (25 – 33°C) and pH (3.5 – 7). The radial growth speed, expressed as colony diameter (cm), under optimal conditions was monitored for 3-8 days (**Figure 1**). Maximum diameter of 8.5 cm minus starting diameter was reached within the time period of 8 days by *R. microsporus, V. volvacea, P. cinnabarinus* and *P. pulmonarius* on day 3, 4, 5 and 8, respectively. The mycelial coverage of *V. volvacea* was relatively thin, but those of the others were white and dense. The slowest radial growth was observed by *F. velutipes, C. comatus* and *S. rugosoannulata*, reaching 0.3, 2.9, 1.3 cm of radial growth at day 4.

**Figure 1.**
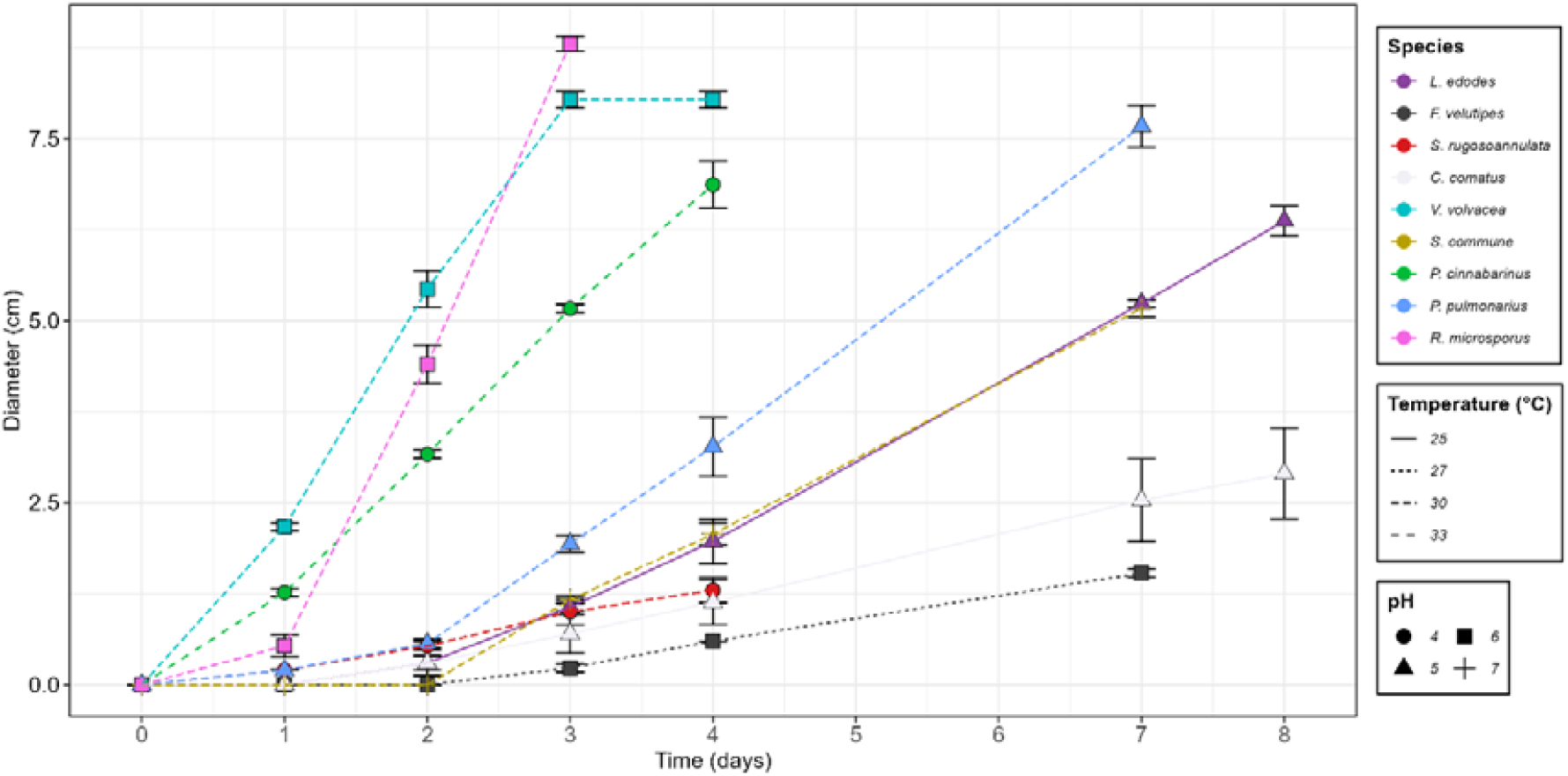
Radial growth of mycelium on malt extract agar plates over time (days) under conditions optimized per species. Color represents the fungal species. Line type represents optimal temperature and point shape represents optimal pH.

Fungal species were cultivated under optimal conditions derived from **Figure 1** in shaking flasks containing malt extract broth. Substrate use, expressed as mycelial biomass yield (%), dry matter loss (%) and residual substrate (%) were monitored (**Figure 2**). Mycelial biomass yield ranged from 10.4% (*S. rugosoannulata*) to 30.7% (*P. pulmonarius*), while *R. microsporus* produced the second highest yield with 27.5%. Dry matter loss described the loss of organic dry matter as volatiles (e.g. CO_2_, NH_3_). The lowest loss in dry weight was observed in *L. edodes,* with 0%. Highest loss in dry matter was observed in *S. commune*, with 68.4%. Residual substrate, substrate that was not oxidized to volatiles, ranged from 9.6% in *P. pulmonarius* to 83.5% in *L. edodes*. Samples with higher yields also had a higher dry matter loss, except *S. rugosoannulata* and *S. commune*, where relatively high dry matter losses were observed with lower yields.

**Figure 2.**
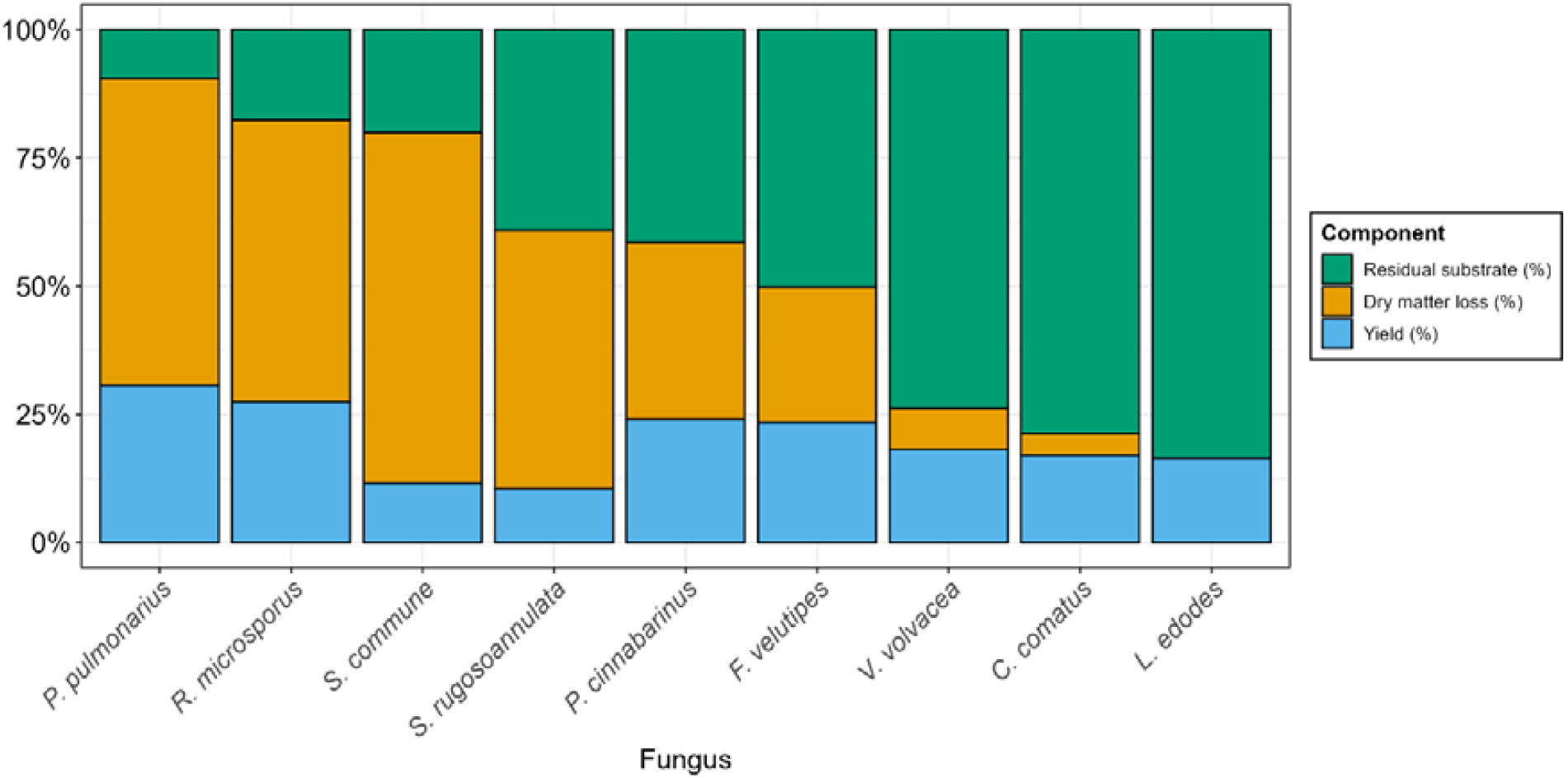
Fungal biomass yield (blue), dry matter loss (orange) and residual substrate (green) of mycelium cultivated in malt extract broth until stationary phase was reached, expressed as percentage of initial substrate dry weight.

### 3.2 Growth dynamics in solid-state fermentation

The 5 most promising species of basidiomycetes were selected for solid-state fermentation (SSF) of brown rice, brewer’s spent grain (BSG) and lupin. Selection criteria were based on growth speed, biomass yield, protein content, protein quality and umami concentration of the mycelium. Overall, the fastest growth speed was observed in *V. volvacea*, the highest yield in *P. pulmonarius*, highest protein content in *S. rugosoannulata*, highest protein quality in *P. cinnabarinus* and highest umami concentration in *S. commune* mycelium. Protein content, protein quality and umami concentration are derived from Zwinkels et al. (2025).

SSF was performed to evaluate the ability of the mycelium to colonize and modify plant-based substrates. Growth was assessed through visual inspection, and mycelial biomass accumulation, was measured through biomarkers glucosamine and ergosterol, while metabolic activity was inferred from changes in pH, temperature profile and changes in substrate composition. The substrates, brown rice, brewer’s spent grain, and lupin, provided distinct nutrient profiles, allowing for comparisons in fungal adaptation and fermentation efficiency.

After incubation, the plant substrates were colonized with mycelium of basidiomycetes and *R. microsporus* (Figure 3). Substrate-fungus combinations that are not depicted did not yield sufficient mycelial biomass accumulation and were therefore not included in further analyses. These pictures provide information on the superficial formation of mycelium. In all samples, mycelia penetrated to the center of the product equally well, relative to their surface mycelial coverage, except for the substrate-fungus combination BSG-*P. cinnabarinus* (Supplementary material 1). In the latter case, the mycelium at the outside was denser than at the inside. All fungi produced a white mycelium, except *V. volvacea*, which had some red pigmented spots. *S. rugosoannulata* and *P. pulmonarius* on BSG, and *S. commune* on brown rice produced a slightly less dense distribution of mycelium on the product, while full and dense mycelial coverage was observed in other fermentations. Out of all tested species of basidiomycetes, only *P. cinnabarinus* could adequately ferment lupin and produce significant amounts of mycelium. Brown rice fermented with *S. commune* mycelium had a softer texture, while brown rice fermented with *V. volvacea* or *P. pulmonarius* had a slightly tougher texture compared to *R. microsporus*. Other basidiomycetous fermentations produced textures similar to the *R. microsporus* fermentations on the same substrate. The fermented samples exhibited distinct aromas depending on the fungal species and substrate. *S. commune* produced a sweet, fruity, and musty aroma. *P. cinnabarinus* had a nutty scent on brown rice, but a strong ammonia smell on BSG. *V. volvacea* produced a mushroom-like odor, *P. pulmonarius* had a pleasant toffee/caramel aroma, and *R. microsporus* produced a clean, subtly sweet scent, similar to fresh tempeh.

**Figure 3.**
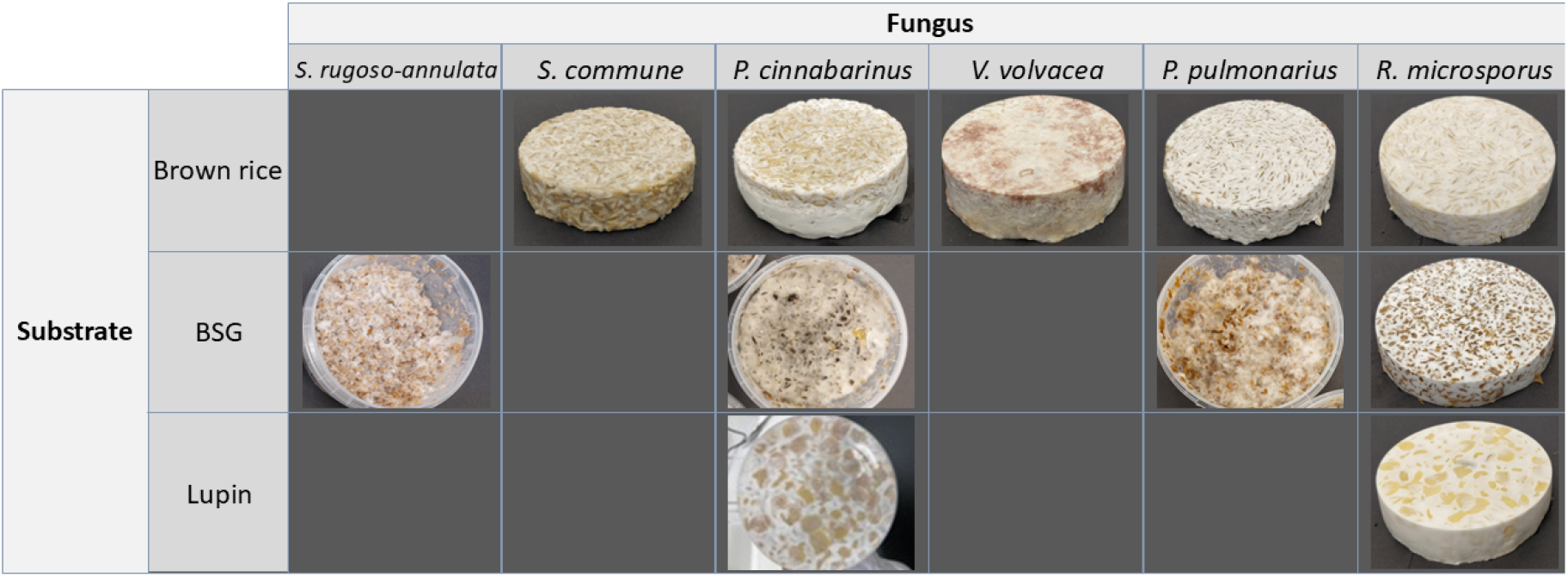
Appearance of brown rice, brewer’s spent grain (BSG) and lupin fermented with *S. rugosoannulata, S. commune, P. cinnabarinus, V. volvacea, P. pulmonarius* and *R. microsporus*.

As indicator for fungal metabolic activity, heat production by the fungi was estimated by monitoring the temperature at the core of samples during the fermentation, and expressed as temperature increase above the incubator temperature (**Figure 4**A). The highest maximum and average temperature increase was observed in *R. microsporus* fermented brown rice, with 6.2°C and 4.0°C, respectively. The core of substrates fermented by *R. microsporus* increased to a stable temperature after around 1 day of fermentation, remaining stable for the rest of the fermentation period. The second highest temperature was observed in substrates fermented by *P. cinnabarinus*, in which temperatures reached their maximum at around day 2, thereafter slowly decreasing. This trend was similarly observed in brown rice – *S. commune*, albeit with a steeper temperature decrease after day 4. Also, the core temperature in fermentations with *V. volvacea* or *P. pulmonarius* increased during the fermentation period. Finally, only a minor temperature increase was observed in *S. rugosoannulata* fermented BSG, with 0.2°C on average. An interesting observation was the circadian rhythm of temperature fluctuations in *P. pulmonarius* fermented BSG, which was repeatedly observed (Supplementary material 2).

**Figure 4.**
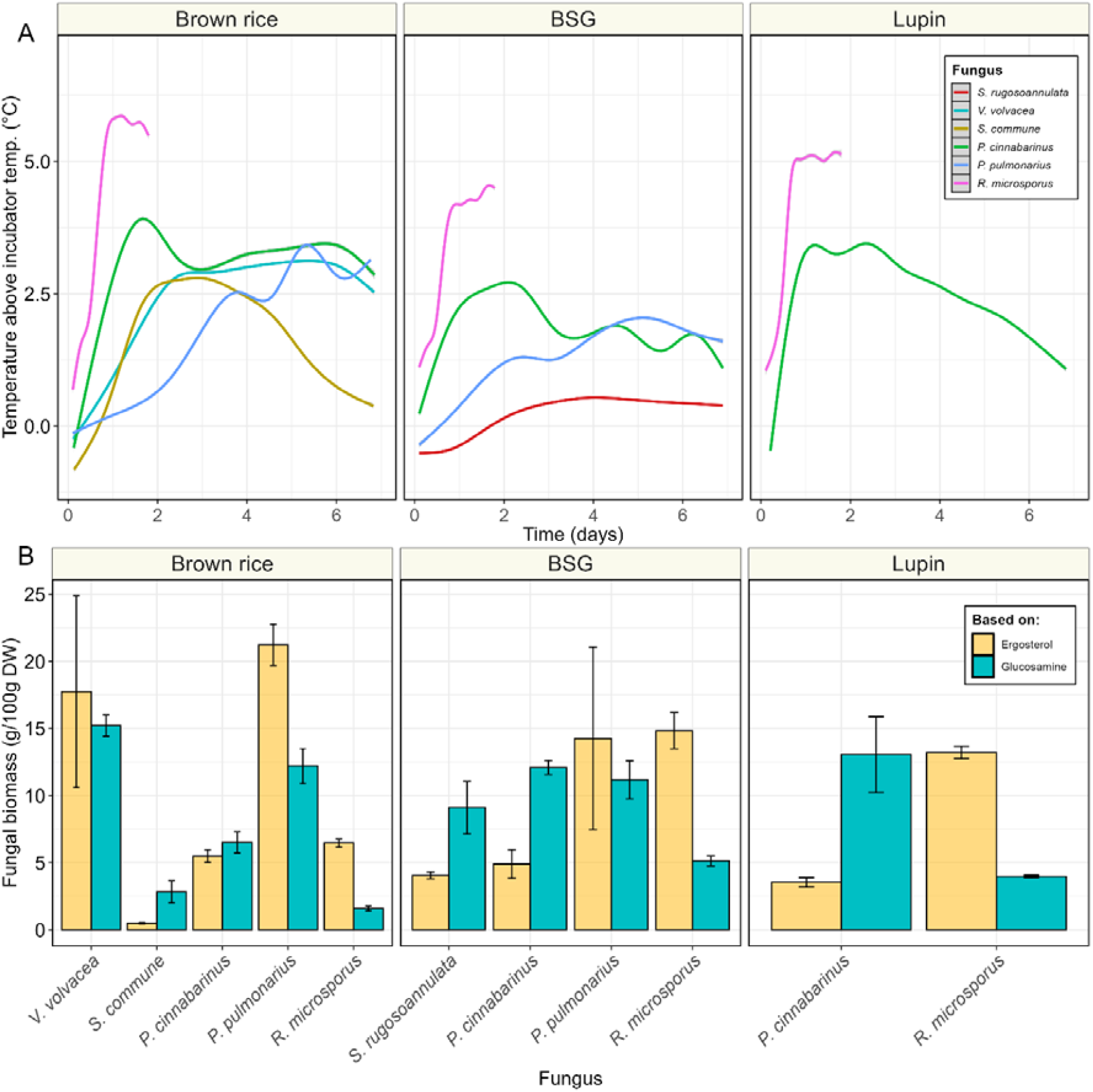
**A)** Temperature profile of substrates (brown rice, BSG & lupin) fermented with *S. rugosoannulata* (black), *S. commune* (orange), *P. cinnabarinus* (green), *V. volvacea* (turquoise), *P. pulmonarius* (blue) or *R. microsporus* (pink). Temperature is indicated as degrees Celsius in the core of the sample above incubator temperature. **B)** Fungal biomass (g/100 g DW) measured through biomarkers ergosterol (yellow) and glucosamine (blue) in fermented samples. Values are means of biological triplicates (± sd).

Besides heat production, fungal biomass (FB) was estimated through biomarkers glucosamine and ergosterol (**Figure 4**B), calibrated based on the concentration found in mycelial biomass per species (Supplementary material 3). Ergosterol based fungal biomass (erg-FB) ranged from 0.5 g/100 g DW in brown rice – *S. commune* to 21.2 g/100 g DW in brown rice – *P. pulmonarius*, while glucosamine based fungal biomass (glu-FB) estimation ranged from 1.6 g/100 g DW in brown rice – *R. microsporus* to 15.2 g/100 g DW in brown rice – *V. volvacea*. Per substrate FB estimations ranged from 0.5 to 21.2 g/100 g DW in brown rice, from 4.0 to 14.8 g/100 g DW in BSG and from 3.5 to 13.2 g/100 g DW in lupin. Erg-FB and glu-FB estimations were largely in agreement in brown rice fermented with basidiomycetes, as well as BSG – *P. pulmonarius*. However, disagreement between the methods was observed in *R. microsporus*, where even though correlation between the methods was high (R^2^ = 0.98; Supplementary material 4) erg-FB estimation was observed to be 2-3 times higher, compared to glu-FB. This discrepancy was inversely observed in BSG, and lupin fermented with *P. cinnabarinus*, which resulted in 1.5-2.5 times higher FB estimations when based on glucosamine.

Overall, *R. microsporus* and *P. cinnabarinus* produced substantial amounts of fungal biomass on all substrates. Brown rice supported the growth of the most tested species of basidiomycetes while also recording the highest temperatures and fungal biomass.

To better understand the fungal metabolic activities in the SSF and how they affect fungal growth and the properties of the fermented substrate, changes in pH, sugars, fibers, amino acids, and dry matter content were monitored. While all substrates were soaked at pH 4.0, pH of unfermented substrates ranged from 5.3 in BSG to 6.7 in brown rice, which is a likely result of differences between the diffusion of acid into the substrate and the buffering capacity of the substrate. Fermentation of brown rice resulted in a minor decrease in pH, strongest in *S. commune*, reaching 4.7 (Figure 5). The pH increased in BSG fermented with *P. cinnabarinus* (to pH 9.0), *P. pulmonarius* (to pH 8.1) and *R. microsporus* (to pH 6.1), as well as *R. microsporus* fermented lupin (to pH 7.8).

**Figure 5.**
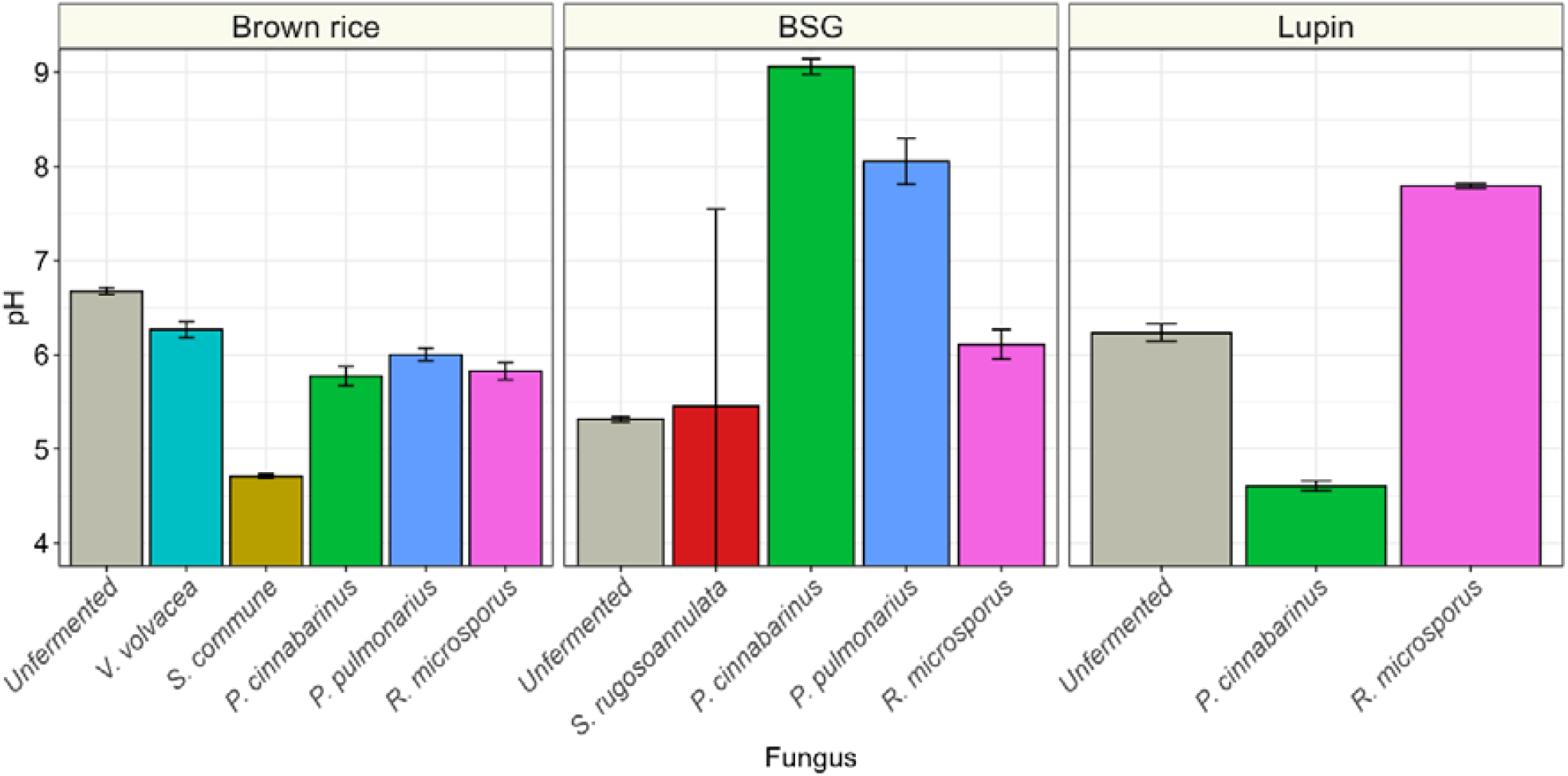
pH values of brown rice, brewer’s spent grain (BSG) and lupin, before fermentation (grey) or fermented with *S. rugosoannulata* (red), *V. volvacea* (turquoise), *S. commune* (orange), *P. cinnabarinus* (green), *P. pulmonarius* (blue) or *R. microsporus* (pink). Values are means of biological triplicates (± sd).

Hydrolysis of starch and hemicellulose result in the formation of neutral sugars, such as glucose and maltose, which are transported into the cytosol via translocators (Fettke et al., 2009). The speed of hydrolysis of polysaccharides to metabolization of simple sugars indicates the suitability of carbohydrate polymers as substrate for basidiomycetes. The high starch content in brown rice, compared to the high hemicellulose/cellulose content in BSG provides insight into the preference of basidiomycetes for these substrates, thereby serving as a model for the fungal hydrolysis activity of carbohydrate polymers. The hydrolysis of complex carbohydrates results in an increased availability of fermentable (simple) sugars like glucose and maltose in all fermentations (**Figure 6**). Concentrations of these compounds were highest in fermented brown rice, with glucose ranging from 0.8 g/100 g DW in *R. microsporus* to 9.7 g/100 g DW in *S. commune* and maltose between 0.4 and 3.5 g/100 g DW in *R. microsporus* and *V. volvacea*, respectively. With glucose and maltose concentrations at 0.1 g/100 g DW in unfermented BSG, increasing to 0.3 – 0.7 g/100 g DW during fermentation, the pool sizes of these free sugars were relatively low in this substrate. On average, basidiomycetous fermentations released more sugars relative to its consumption than *R. microsporus*-based fermentations.

**Figure 6.**
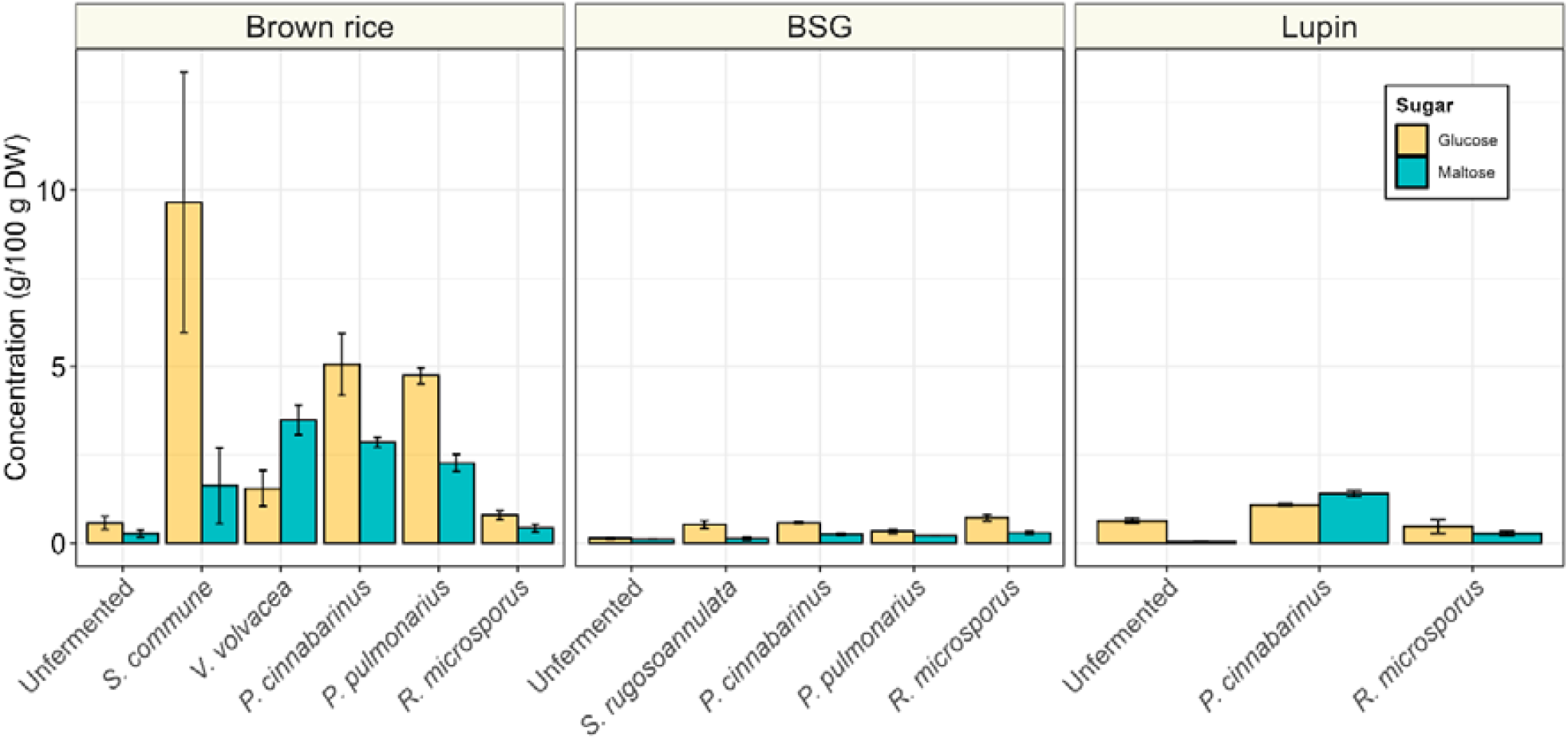
Glucose (yellow) and maltose (blue) concentrations (g/100 g DW) of brown rice, brewer’s spent grain (BSG) and lupin, before fermentation or fermented with *S. rugosoannulata, S. commune, P. cinnabarinus, V. volvacea, P. pulmonarius* or *R. microsporus*. Values are means of biological triplicates (± sd).

BSG, the industry side-stream of beer brewing, has a low sugar and starch concentration, due to the malting and mashing process (Briggs et al., 1981). This enzymatic process liberates amylolytic enzymes, which break down starch to mainly maltose. Thereafter, this maltose gets rinsed out during the lautering step. Basidiomycetes are known to degrade complex carbohydrates, such as lignin and cellulose, which are generally high in BSG. Therefore, changes in the fiber content of BSG during fermentation were estimated through acid detergent fiber (ADF) and neutral detergent fiber (NDF) content (**Figure 7**). The ADF fraction is largely composed of cellulose and lignin (Van Soest, 1973), while NDF also contains hemicellulose (Van Soest et al., 1991). The ADF and NDF content in unfermented BSG was 21.2 and 62.6 g/100 g DW, respectively. ADF did not significantly change during fermentation, while NDF only decreased in *S. rugosoannulata* to 57.4 g/100 g DW (p > 0.05) and in *P. cinnabarinus* to 47.4 g/100 g DW (*p* < 0.05), thereby indicating that this decrease was mostly a result of hemicellulose degradation. The absence of fiber degradation in *R. microsporus* was confirmed by assessment of the fiber content based on the original dry weight, before fermentation (**Figure 7**B). When considering the higher loss in dry weight in basidiomycetes, compared to *R. microsporus*, a loss in NDF and mainly ADF was observed in basidiomycetes, but remained absent in *R. microsporus* fermentations.

**Figure 7.**
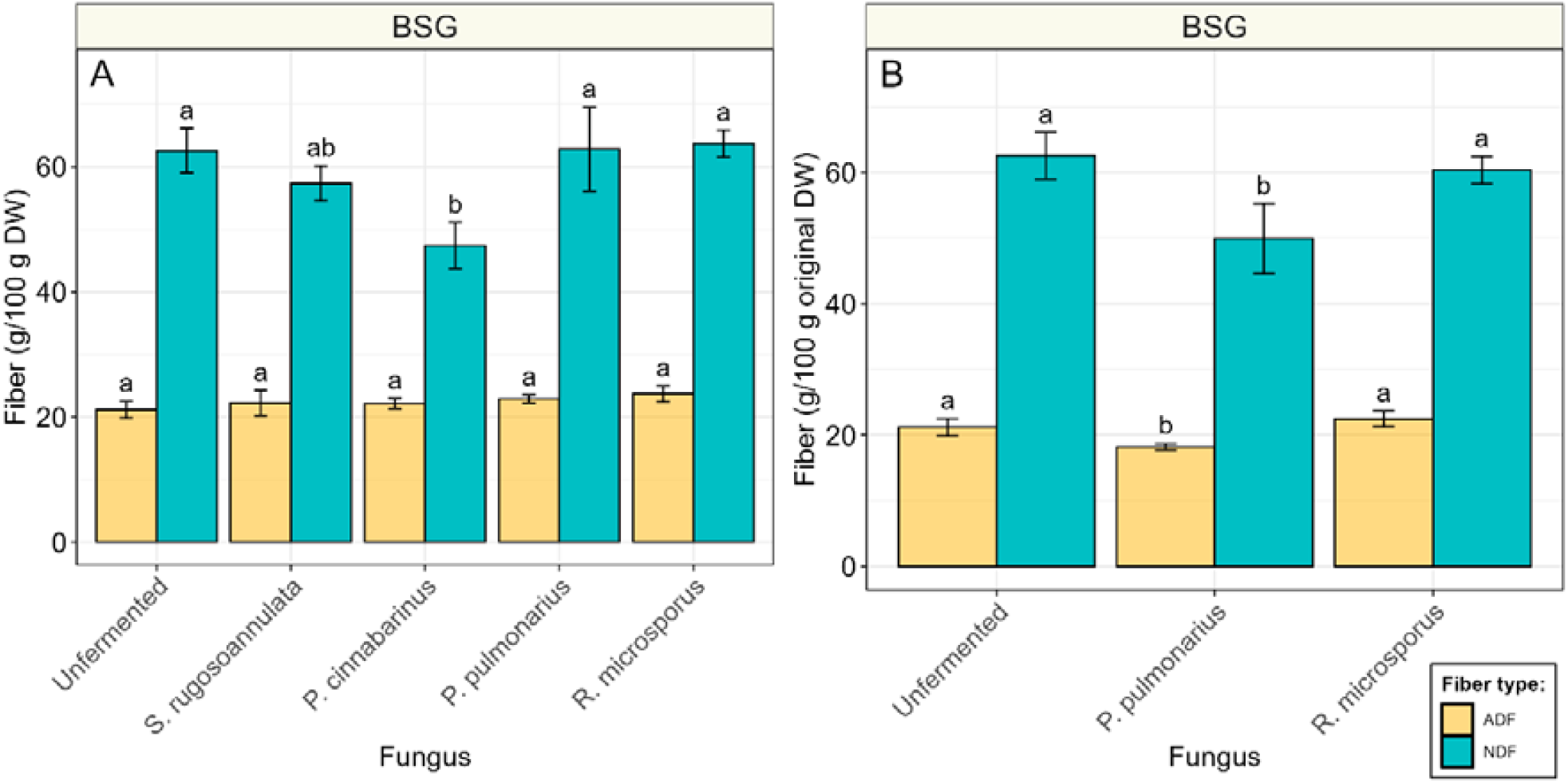
Acid detergent fiber (ADF, yellow) and neutral detergent fiber (NDF, blue) concentrations of brewer’s spent grain (BSG), before fermentation and fermented with *S. rugosoannulata, P. cinnabarinus, P. pulmonarius* or *R. microsporus*. **A)** Fiber content based on dry weight. **B)** Fiber content of subset based on content of dry weight of unfermented sample. Values are means of biological triplicates (± sd). Different letters indicate significant differences (*p* < 0.05).

In addition to carbohydrate metabolism, nitrogen metabolism was investigated. We monitored the changes in dry matter loss, crude protein content, and total amino acid content during fermentation (Figure 8A and 8B). An increase in dry matter loss reflects the reduction of dry matter compared to the unfermented substrate. If nitrogen and amino acid levels changed to the same extend, this would imply that the observed changes are solely due to dry matter loss. However, deviations from this pattern suggest either nitrogen loss through volatilization (in the case of crude protein) or amino acid degradation (in the case of total amino acids).

**Figure 8.**
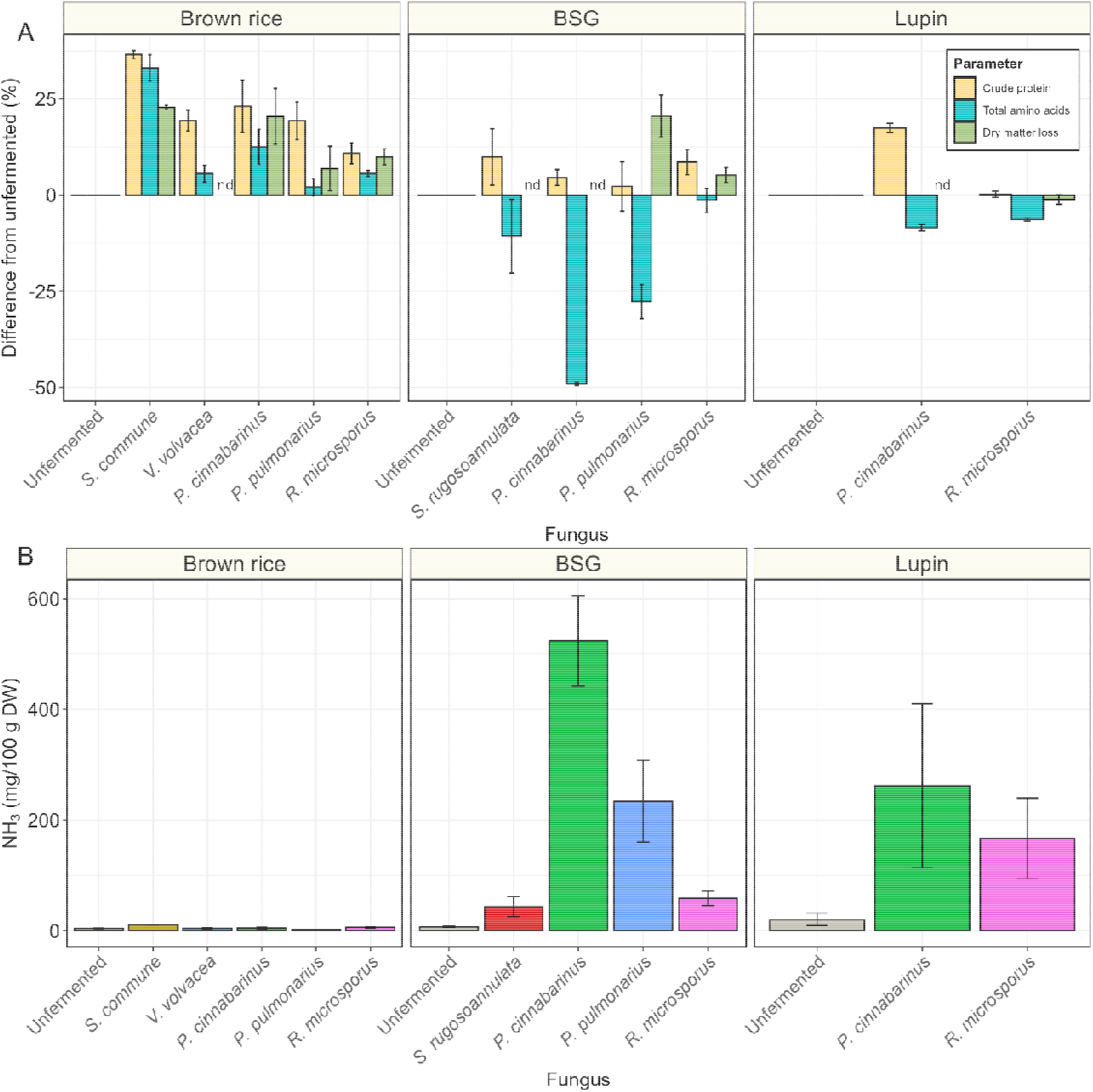
**A)** Relative change of crude protein (yellow), total amino acids (blue) and dry matter loss (green) after fermentation, expressed at percentage change from the unfermented substrate (brown rice, brewer’s spent grain (BSG) and lupin). Nd = not determined. **B)** Ammonia content (mg/100 g DW) of substrates unfermented (grey) or fermented with *S. rugosoannulata* (red), *S. commune* (orange), *P. cinnabarinus* (green), *V. volvacea* (turquoise), *P. pulmonarius* (blue) or *R. microsporus* (pink). Values are means of biological triplicates (± sd).

In brown rice, fermentation with basidiomycetes and *R. microsporus* resulted in a proportional increase in crude protein, total amino acids and dry matter. The is observed as an increase in crude protein and amino acid content up to 36.5% and 33.0%, respectively. However, in BSG and lupin fermentations the changes in these parameters were not proportional. Particularly, in BSG an increase in crude protein and loss in dry matter were matched by a decrease in total amino acids. The loss in total amino acids corresponds well with the ammonia produced during fermentation (**Figure 8**B). The loss of amino nitrogen, combined with the increase in total nitrogen, and the formation of ammonia indicate the occurrence of both proteolysis and deamination in these substrates.

A major benefit of fungal fermentation is the degradation of anti-nutritional factors (ANFs) (Samtiya et al., 2020). ANFs are often high in grains and legumes and have been reported to reduce mineral absorption and protein digestion (Samtiya et al., 2020). A major ANF with these properties in plant foods is phytic acid (Gibson et al., 2018). Phytic acid was found to be highest in unfermented BSG (2.9 g/100 g DW), followed by unfermented brown rice (1.2 g/100 g DW) and unfermented lupin (0.56 g/100 g DW) (**Figure 9**). Fermentation with all of the selected species of basidiomycetes resulted in a significant reduction of phytic acid of 69-80% in brown rice, 63-78% in BSG, and 74% in lupin. In contrast, phytic acid was not significantly reduced in lupin and brown rice fermented with *R. microsporus* and phytic acid reduction was significantly lower for *R. microsporus* in BSG (23% reduction). These findings suggest that phytic acid is particularly well degraded by basidiomycetes.

**Figure 9.**
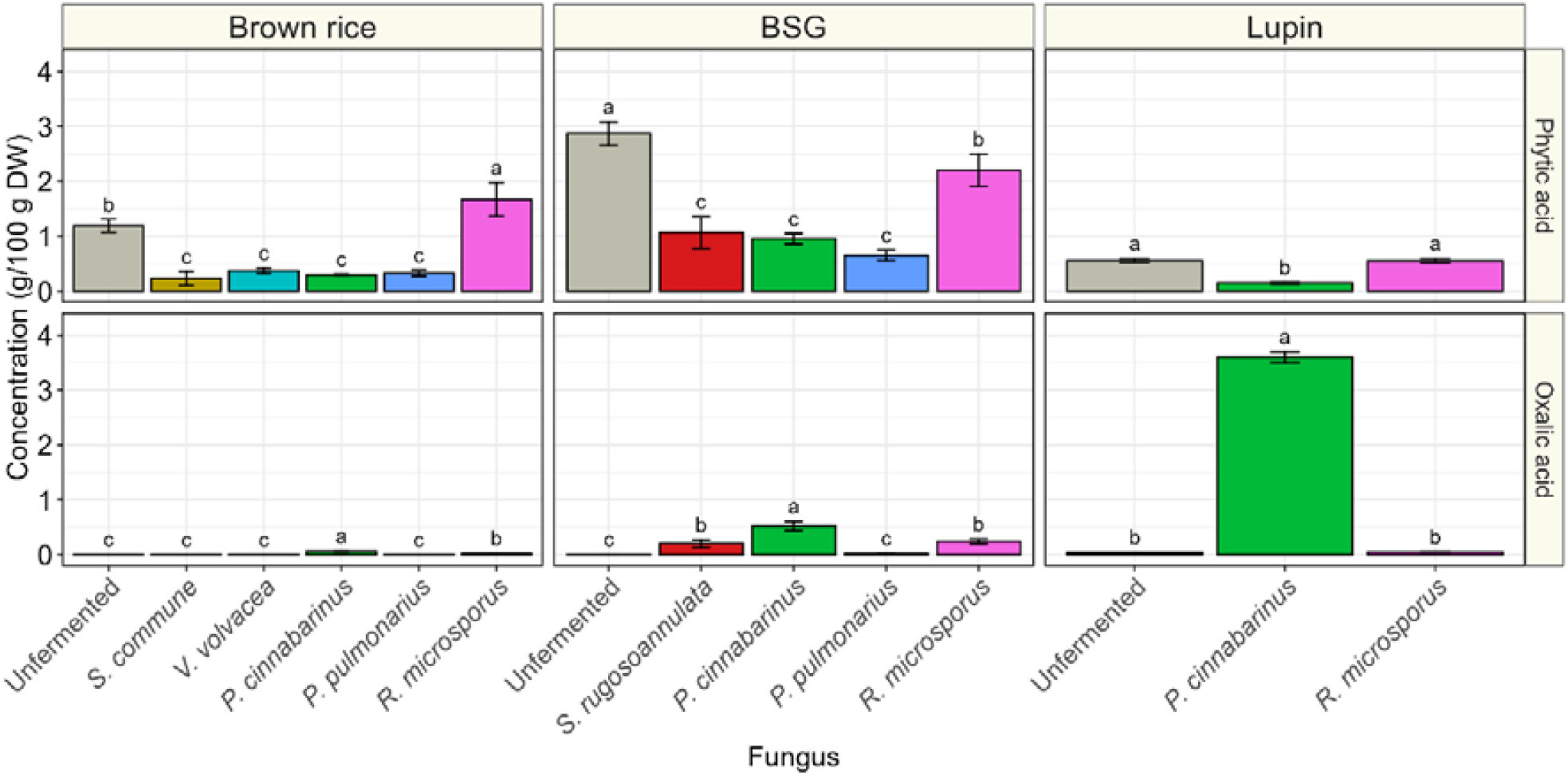
Oxalic acid and phytic acid concentration (g/100 g DW) of unfermented (grey) substrates, brown rice, brewer’s spent grain (BSG) and lupin, or substrates fermented with *S. rugosoannulata* (red), *S. commune* (orange), *P. cinnabarinus* (green), *V. volvacea* (turquoise), *P. pulmonarius* (blue) or *R. microsporus* (pink). Values are means of biological triplicates (± sd). Different letters indicate significant differences (based on comparisons within substrate; *p* < 0.05).

Moreover, oxalic acid is an another ANF that is found in plants, but can also be produced by fungi, and particularly by basidiomycetes, where oxalic acid is typically one of the major organic acids produced (Galkin et al., 1998). Like phytic acid, also oxalic acid chelates minerals, thereby reducing their bioavailability (Noonan, 1999). In plants, it is particularly high in rhubarb (3.3-9.5 g/100 g DW), and spinach (7-14 g/100 g DW), but concentrations are relatively low in cereals (Kawazu et al., 2003; Libert & Creed, 1985). Oxalic acid concentrations in unfermented substrates were 8.2, 8.4, and 42 mg/100 g DW, in brown rice, BSG and lupin, respectively (**Figure 9**). Oxalic acid concentrations were particularly increased in substrates fermented by *P. cinnabarinus*, which resulted in 0.059 g/100 g DW in brown rice (p>0.05), 0.52 g/100 g DW in BSG (p < 0.05), and 3.6 g/100 g DW in lupin (p < 0.05). In addition, oxalic acid significantly increased in BSG fermented with *S. rugosoannulata* or *R. microsporus* to 0.20 and 0.24 g/100 g DW, respectively. Therefore, multiple fungi can produce oxalic acid, but the extent is substrate and species dependent.

## 4 Discussion

This study explored the mycelial growth dynamics and metabolic activity of a selection of species of basidiomycetes in liquid and solid-state fermentations, assessing their impact on substrate composition and nutritional parameters. By optimizing growth conditions and analyzing biochemical changes, we aimed to understand how these fungi modify food-grade, plant-based substrates. Here, we discuss the key findings in relation to fungal growth, metabolism, and their potential for improving nutritional properties.

Growth speed is a critical factor in solid-state fermentation (SSF), as industrial applications require short fermentation periods to become economically viable. Basidiomycetes naturally colonize lignocellulosic materials such as straw and wood, but their adaptation to these nutritionally poor substrates often results in slow growth on these substrates, with full colonization and fruiting body formation taking months. To make solid-state fermentation commercially feasible, optimizing environmental conditions to accelerate fungal growth is essential. Among the tested fungi, only *V. volvacea* and *P. cinnabarinus* matched the growth speed of *R. microsporus* on agar plates (**Figure 1**). Notably, *P. cinnabarinus* exhibited rapid growth at pH 4, making it particularly promising for industrial fermentations, since the low pH reduces or even prevents growth of many spoilage and pathogenic bacteria (Nout et al., 1987).

In addition to rapid growth, high substrate conversion efficiency is a key factor in selecting species for solid-state fermentation (SSF). To assess this, we determined fungal biomass yields and dry matter loss in submerged fermentations (**Figure 2**). The highest combined yield and dry matter loss, representing total substrate utilization, was observed in *P. pulmonarius*. This species did not grow the fastest on agar plates, suggesting that metabolic strategies for fast colony expansion on agar plates is not a good predictor for effective substrate utilization in a solid-state fermentation. The high yields observed in *P. pulmonarius* and *P. cinnabarinus* (both belonging to the order of Polyporales) suggest a potential metabolic advantage within this order promoting efficient substrate conversion. Conversely, the high dry matter loss relative to fungal biomass yield in *S. rugosoannulata* and *S. commune* indicates that these species likely had a more inefficient ATP to CO_2_/H_2_O ratio or a higher maintenance requirement. *S. commune* particularly has a low biomass yield and high dry matter loss, both in submerged fermentation and in solid-state fermentation (SSF). Combined with the high glucose yield on brown rice, *S. commune* also lowered the pH which indicates acetic acid secretion, which has a high CO_2_-ATP ratio.

Fungal biomass estimation in SSF is more complex than in submerged fermentation due to the intertwined nature of the mycelium and the substrate, making direct fungal biomass measurement impossible (Steudler & Bley, 2015a). To address this, we employed a combination of indirect methods: visual inspection, heat production, and biomarker analysis using ergosterol and glucosamine. Each of these methods has inherent limitations, such as differentiation between types of mycelia, superficial estimation, and requirement of careful (*ex situ*) calibration. Using a combination of these methods diminishes these limitations, allowing for the estimation of different types of fungal biomass, e.g. inactive/active, dead/alive, and verification.

Generally, fungal biomass estimation provided a complete overview of the formation of fungal biomass. The methods based on biomarkers, ergosterol and glucosamine, were largely in agreement, and discrepancies can be diminished by the combination of methods. By combining different fungal biomass estimation methods, we found that the selected basidiomycete species generally exhibited lower metabolic activity compared to *R. microsporus*, and required more specific nutritional conditions, as reflected by the selective growth behaviour of some species. However, with a slightly prolonged fermentation time, also these fungi can completely colonize the high-starch cereal brown rice, the low-starch cereal BSG and the high-protein legumes lupin (**Table 2**). Adequate biomass formation is imperative for selection of the right basidiomycetous species. This was clearly demonstrated by the observed large variation in success rate and biomass yield (**Figure 3**).

In addition to fungal biomass formation, alterations in the chemical composition of the substrates were assessed to determine differences in metabolic behavior of the basidiomycetes and *R. microsporus* on substrates with distinct nutritional compositions. The high starch content in brown rice was an easily accessible substrate for the basidiomycetes and was likely the main carbon source used. The preference for and degradation of starch is indicated by the high glucose and maltose concentrations in basidiomycete-fermented brown rice, which suggests enzymatic hydrolysis of starch at higher rates than uptake and utilization of glucose by the fungal mycelium. Secondly, the decrease in pH in brown rice likely originates from glucose degradation, resulting in the formation of organic acids. The accessibility of starch as a carbon source is evidenced by the rapid initial temperature rise and final high sugar content in brown rice, in contrast to the low-starch substrates, BSG and lupin.

Interestingly, also low-starch, high-protein substrates, like BSG and lupin, supported the formation of substantial amounts of fungal biomass for some of the selected basidiomycetes, despite the lower apparent starch degradation. Moreover, fiber degradation in BSG seemed to only play a minor role in energy production, despite the well-known ability of basidiomycetes to grow on lignocellulosic material (Briggs et al., 1981; Lundell et al., 2010). However, this lignocellulosic material is generally low in nitrogen (0.04-0.59%) (Martin et al., 2014; van hecke 2020)). The higher (protein-derived) nitrogen content in BSG and lupin seemed to be a more accessible substrate than fibers. This process led to the deamination of amino acids, the production of ammonia and thereby an increase in pH observed in these fermented substrates. Only in *P. cinnabarinus* fermented lupin, a decrease in pH was observed, which is linked to the high oxalic acid production. These results indicate that basidiomycetes prioritize fast growth in nutrient rich substrates by deamination of more easily accessible amino acids over slower, but more efficient growth on fibers. The loss of amino acids results in a reduction of the nutritional value due to this fermentation. To prevent this, the presence of an easily metabolizable carbon source in the substrate, as is present in brown rice, appears critical for preserving amino acids and, consequently, nutritional quality. This effect is most evident in basidiomycetes, but the modest decline in total amino acids and simultaneous rise in ammonia observed in *R. microsporus*-fermented BSG and lupin similarly support this concept. Lastly, the discrepancies between the crude protein and amino acid content in fermented BSG and lupin could lead to an overestimation of actual protein by the crude protein method, since the deamination of amino acids did not decrease the nitrogen content to the same extent. Therefore, DUMAS and Kjeldahl methods could overestimate the protein content in fungal fermented substrates. Therefore, measuring total amino acids is proposed as a more accurate approach for assessing the protein content of fungal-fermented substrates.

Fungal fermentation can influence anti-nutritional factors (ANFs) in plant-based substrates as well. These reduction strategies gain importance in high plant-based diets, which are more prone to lower mineral intake and absorption (Neufingerl & Eilander, 2022). Fungi, and in particular basidiomycetes are well-known to produce a range of enzymes degrading ANFs, such as phytases, thereby improving the nutritional value of a food product (J. Wang et al., 2023). A major ANF in cereals is phytic acid, which is known to reduce mineral bioavailability and hinder protein digestion. High intake of phytic acid, combined with low mineral and protein intake can lead to deficiencies in these nutrients (Dahdouh et al., 2019; Gibson et al., 2018; Lopez et al., 2002). For this reason, phytic acid content of BSG has been viewed as a major hurdle for inclusion as ingredient in food products (Lynch et al., 2016). The phytic acid content in our samples of the unfermented brown rice was found to be _∼_1.0 g/100 g DW), which is in the range of what is reported in literature (Liang et al., 2007). On the other hand, we found slightly higher phytic acid concentrations in BSG and lupin, compared to literature, namely 2.9, and 0.56 g/100 g DW in BSG and lupin, respectively, instead of 1.0 g/100 g DW and 0.063% (Ktenioudaki et al., 2015; Prusinski, 2017). The significant reduction in phytic acid in all substrates of up to 80% suggest that basidiomycetes are well equipped to produce phytases in SSF, as has been observed earlier by da Luz et al. (2013) in Jatropha curcas L. seed cake fermented with Pleurotus ostreatus. In *R. microsporus*-based fermentations the degradation of phytic acid was either less or absent. The ability of *R. microsporus* to produce phytases has been documented, so the absence of reduction is likely due to low expression of this enzyme (Azeke et al., 2011). A substrate-dependent phytase expression has been observed before. For instance, a 30% reduction was observed in fermented soybeans (Egounlety & Aworh, 2003), whereas no significant reduction was found in fermented BSG biscuits (X. Wang et al., 2023).

In contrast to phytic acid, oxalic acid is an ANF that is often produced by white-rot fungi (Galkin et al., 1998), originating from intermediates of the tricarboxylic acid (TCA) cycle and serving a function by promoting manganese-peroxidase activity, which breaks down lignin (Dutton & Evans, 2011; Grąz, 2024). Therefore, the high oxalic acid content in *P. cinnabarinus*-based fermentations could be a result of higher activity of ligninolytic enzymes in this sample, which corresponds with the higher degradation of acid detergent fiber (ADF) and its well-documented ability to produce ligninolytic enzymes (Levasseur et al., 2014). The anti-nutrient effect of oxalic acid stems from its capacity to chelate minerals, such as calcium, iron, and zinc, reducing their bioavailability, which can lead to calcium deficiencies in people with diets high in oxalate and low in calcium (Dagostin, 2016; Noonan, 1999). In SSF, oxalic acid production seemed to be dependent on the combination of substrate and fungal species. The ANF was mainly produced in *P. cinnabarinus* fermentations, which was highest in lupin. The oxalic acid content of lupin-*P. cinnabarinus* (3.6 g/100 g DW) exceeds that of other high-oxalate legumes, such as white beans (0.5 g/100 g dry beans), soybeans (0.3 g/100 g dry beans), but is still lower than high-oxalate foods, such as rhubarb (3.3-9.5 g/100 g DW), and spinach (7-14 g/100 g DW) (Kawazu et al., 2003; Libert & Creed, 1985). The nutritional benefit of SSF with basidiomycetes depends on substrate and species formulation. While phytic acid was universally reduced, oxalic acid production was both species and substrate dependent.

Overall, these results show that basidiomycetes can fully colonize cereals and legumes, producing high fungal biomass and dense mycelial coverage, but at a slower rate than *R. microsporus*. The distinct nutritional compositions of the substrates, serving as a model for high-starch, or high-protein plant-foods, respectively, resulted in similar growth patterns across the selected species of basidiomycetes. The pattern of deamination in high-protein substrates, combined with its absence in the high-starch brown rice suggest the requirement of a readily available carbon source to prevent loss of protein. When substrate composition and fungal species were optimally matched, this resulted in high mycelial yields, increased protein concentration, and reduction of phytic acid concentrations. Moreover, the variability in growth rate and fungal biomass formation (“yield”) between the selected species of basidiomycetes, indicates the potential of further optimization of biomass formation. The general agreement of radial growth data and biomass yield data in submerged fermentation with biomass formation in solid-state fermentation suggest that radial growth could serve as a practical screening metric for selecting basidiomycetes suitable for solid-state fermentation of plant foods. Future studies will explore the nutritional and sensory advantages, particularly regarding protein quality and umami flavor, of basidiomycete-based fermentations compared to traditional fungal fermentations.

## 5 Conclusion

This study demonstrates the potential of a number of species of basidiomycetes in solid-state fermentation of plant-based substrates, by assessing a phylogenetically heterogenous selection of species on diverse substrates. Substrate formulation plays a crucial role in fungal growth and metabolism. An easily accessible carbon source, such as starch is required in high-protein substrates to prevent the catabolism of amino acids. With the correct formulation, species belonging to the phylum of Basidiomycota can form a dense mycelial network comparable to that found in tempeh. This food fermentation process leads to an increase protein content and degradation of anti-nutritional factors, such as phytic acid. The latter process is more effective as compared to conventional solid-state fermentation processes driven by *Rhizopus* sp. The methodology applied in this study can serve as a selection tool for the expansion of basidiomycete species in solid-state food fermentation processes.

## Supporting information

Supplementary file Zwinkels et al. 2025

## CRediT authorship contribution statement

**Jasper Zwinkels:** Conceptualization, Methodology, Investigation, Formal analysis, Data curation, Validation, Writing – original draft. **Stef van Oorschot**: Investigation, Formal analysis. **Oscar van Mastrigt**: Conceptualization, Methodology, Writing – review & editing, Supervision. **Eddy J. Smid**: Conceptualization, Methodology, Writing – review & editing, Supervision, Funding acquisition, Project administration.

## Acknowledgements

We would like to thank Philip van Koolwijk for his contribution to the production and analysis of mycelial biomass, Ran Xu and Alessia de Chaud for their exploratory work into solid-state fermentation using basidiomycetes, and Dr. Arend van Peer and Dr. Karin Scholtmeijer for their expert opinion on basidiomycete selection. We gratefully acknowledge Dr. Rebecca Rocchi for her inspiration in the graphical design. Lastly, we would like to thank Dr. Nikkie van der Wielen and Dr. Leon de Jonge for their advice and performing the amino acid extraction and analysis.

## Data availability

Data will be made available on request.

