## Supplementary file Zwinkels et al. 2025 for "The potential of mycelium from mushroom-producing fungi in alternative protein production: a focus on fungal growth, metabolism, and nutrition"

### 1 Supplementary material

**Supplementary material 1 |** Cross-sectional appearance of brown rice, brewer's spent grain (BSG) and lupin fermented with *S. rugosoannulata*, *S. commune, P. cinnabarinus*, *V. volvacea*, *P. pulmonarius* and *R. microsporus*.

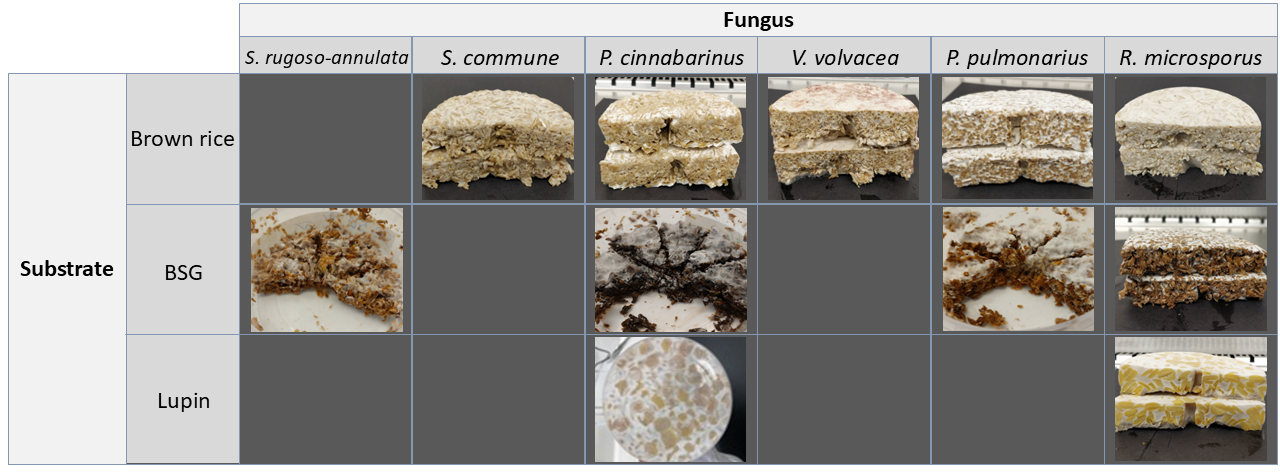

**Supplementary material 3 |** Temperature profile of Brewer’s spent grain fermented by *P. pulmonarius* at different maltose concentration (A; 0-10%) or pH values (B;3.5-6.5).

Temperature (°C)

| Fermentation time (days) | 0% | 1% | 3% | 5% | 8% | 10% |
| --- | --- | --- | --- | --- | --- | --- |
| 0.222222 | 28.812 | 28.562 | 28.688 | 28.688 | 28.625 | 28.812 |
| 0.229167 | 28.875 | 28.688 | 28.812 | 28.75 | 28.75 | 28.875 |
| 0.236111 | 28.938 | 28.75 | 28.875 | 28.875 | 28.812 | 28.938 |
| 0.243056 | 29 | 28.875 | 28.938 | 28.938 | 28.875 | 28.938 |
| 0.25 | 29.062 | 28.875 | 29 | 29 | 28.938 | 28.938 |
| 0.256944 | 29.062 | 28.938 | 29.062 | 28.938 | 28.938 | 29 |
| 0.263889 | 29.062 | 28.938 | 29.062 | 29 | 29 | 29 |
| 0.270833 | 29.125 | 29 | 29.062 | 29 | 29 | 29.062 |
| 0.277778 | 29.125 | 29 | 29.125 | 29.062 | 29 | 29 |
| 0.284722 | 29.062 | 29.062 | 29.062 | 29 | 29.062 | 29 |
| 0.291667 | 29.125 | 29 | 29.125 | 29 | 29 | 29 |
| 0.298611 | 29.062 | 29 | 29.062 | 29 | 29.062 | 28.938 |
| 0.305556 | 29.125 | 29 | 29.062 | 29 | 29 | 28.938 |
| 0.3125 | 29.062 | 29 | 29.062 | 29 | 29 | 28.938 |
| 0.319444 | 29.062 | 29 | 29.062 | 29 | 29 | 28.938 |
| 0.326389 | 29.062 | 29 | 29.062 | 29 | 29 | 29 |
| 0.333333 | 29.062 | 29 | 29.062 | 28.938 | 29.062 | 28.938 |
| 0.340278 | 29.062 | 29.062 | 29.062 | 29 | 29.062 | 29 |
| 0.347222 | 29.062 | 29 | 29.062 | 28.938 | 29.062 | 28.938 |
| 0.354167 | 29.062 | 29 | 29.062 | 28.938 | 29.062 | 29 |
| 0.361111 | 29.062 | 29 | 29.062 | 29 | 29 | 29 |
| 0.368056 | 29.125 | 29.062 | 29.062 | 29 | 29.125 | 29 |
| 0.375 | 29.125 | 29.062 | 29.125 | 29 | 29.062 | 29 |
| 0.381944 | 29.062 | 29.062 | 29.125 | 29.062 | 29.062 | 29 |
| 0.388889 | 29.125 | 29.062 | 29.188 | 29 | 29.062 | 29.062 |
| 0.395833 | 29.125 | 29.125 | 29.188 | 29.062 | 29.125 | 29.062 |
| 0.402778 | 29.125 | 29.125 | 29.188 | 29.062 | 29.125 | 29.125 |
| 0.409722 | 29.188 | 29.125 | 29.188 | 29.062 | 29.125 | 29.125 |
| 0.416667 | 29.125 | 29.188 | 29.25 | 29.125 | 29.188 | 29.125 |
| 0.423611 | 29.188 | 29.125 | 29.188 | 29.125 | 29.188 | 29.125 |
| 0.430556 | 29.188 | 29.188 | 29.188 | 29.125 | 29.188 | 29.125 |
| 0.4375 | 29.25 | 29.188 | 29.188 | 29.125 | 29.188 | 29.125 |
| 0.444444 | 29.25 | 29.125 | 29.25 | 29.125 | 29.188 | 29.125 |
| 0.451389 | 29.25 | 29.188 | 29.188 | 29.188 | 29.188 | 29.188 |
| 0.458333 | 29.25 | 29.25 | 29.25 | 29.188 | 29.25 | 29.188 |
| 0.465278 | 29.25 | 29.25 | 29.25 | 29.188 | 29.25 | 29.188 |
| 0.472222 | 29.25 | 29.25 | 29.312 | 29.188 | 29.25 | 29.188 |
| 0.479167 | 29.312 | 29.25 | 29.312 | 29.188 | 29.312 | 29.188 |
| 0.486111 | 29.312 | 29.312 | 29.312 | 29.25 | 29.25 | 29.25 |
| 0.493056 | 29.312 | 29.312 | 29.312 | 29.25 | 29.312 | 29.25 |
| 0.5 | 29.312 | 29.312 | 29.375 | 29.25 | 29.312 | 29.312 |
| 0.506944 | 29.375 | 29.312 | 29.375 | 29.312 | 29.375 | 29.375 |
| 0.513889 | 29.375 | 29.312 | 29.375 | 29.375 | 29.375 | 29.312 |
| 0.520833 | 29.375 | 29.375 | 29.438 | 29.312 | 29.375 | 29.375 |
| 0.527778 | 29.375 | 29.438 | 29.438 | 29.375 | 29.438 | 29.375 |
| 0.534722 | 29.438 | 29.438 | 29.5 | 29.438 | 29.438 | 29.438 |
| 0.541667 | 29.438 | 29.438 | 29.5 | 29.375 | 29.438 | 29.375 |
| 0.548611 | 29.438 | 29.438 | 29.5 | 29.438 | 29.438 | 29.438 |
| 0.555556 | 29.5 | 29.5 | 29.5 | 29.5 | 29.438 | 29.438 |
| 0.5625 | 29.5 | 29.5 | 29.562 | 29.438 | 29.5 | 29.438 |
| 0.569444 | 29.562 | 29.562 | 29.562 | 29.5 | 29.5 | 29.5 |
| 0.576389 | 29.5 | 29.562 | 29.625 | 29.5 | 29.562 | 29.5 |
| 0.583333 | 29.562 | 29.562 | 29.562 | 29.5 | 29.562 | 29.5 |
| 0.590278 | 29.562 | 29.562 | 29.688 | 29.5 | 29.562 | 29.562 |
| 0.597222 | 29.625 | 29.625 | 29.625 | 29.562 | 29.562 | 29.562 |
| 0.604167 | 29.625 | 29.625 | 29.625 | 29.562 | 29.625 | 29.562 |
| 0.611111 | 29.625 | 29.625 | 29.688 | 29.562 | 29.625 | 29.625 |
| 0.618056 | 29.688 | 29.625 | 29.75 | 29.625 | 29.688 | 29.625 |
| 0.625 | 29.688 | 29.688 | 29.75 | 29.688 | 29.688 | 29.625 |
| 0.631944 | 29.688 | 29.625 | 29.75 | 29.688 | 29.688 | 29.625 |
| 0.638889 | 29.75 | 29.688 | 29.75 | 29.625 | 29.688 | 29.688 |
| 0.645833 | 29.688 | 29.75 | 29.812 | 29.688 | 29.75 | 29.688 |
| 0.652778 | 29.75 | 29.75 | 29.812 | 29.75 | 29.75 | 29.688 |
| 0.659722 | 29.75 | 29.812 | 29.812 | 29.688 | 29.75 | 29.688 |
| 0.666667 | 29.75 | 29.75 | 29.812 | 29.75 | 29.75 | 29.75 |
| 0.673611 | 29.812 | 29.75 | 29.875 | 29.75 | 29.812 | 29.75 |
| 0.680556 | 29.812 | 29.812 | 29.875 | 29.75 | 29.812 | 29.75 |
| 0.6875 | 29.812 | 29.75 | 29.875 | 29.812 | 29.75 | 29.688 |
| 0.694444 | 29.812 | 29.812 | 29.875 | 29.812 | 29.812 | 29.75 |
| 0.701389 | 29.812 | 29.812 | 29.875 | 29.812 | 29.812 | 29.75 |
| 0.708333 | 29.812 | 29.875 | 29.875 | 29.812 | 29.812 | 29.812 |
| 0.715278 | 29.812 | 29.812 | 29.875 | 29.812 | 29.812 | 29.75 |
| 0.722222 | 29.875 | 29.812 | 29.875 | 29.812 | 29.875 | 29.75 |
| 0.729167 | 29.875 | 29.812 | 29.938 | 29.812 | 29.875 | 29.812 |
| 0.736111 | 29.875 | 29.812 | 29.875 | 29.812 | 29.875 | 29.812 |
| 0.743056 | 29.875 | 29.875 | 29.938 | 29.812 | 29.875 | 29.812 |
| 0.75 | 29.875 | 29.875 | 29.938 | 29.812 | 29.875 | 29.812 |
| 0.756944 | 29.875 | 29.938 | 29.938 | 29.875 | 29.875 | 29.812 |
| 0.763889 | 29.875 | 29.875 | 29.938 | 29.875 | 29.875 | 29.812 |
| 0.770833 | 29.875 | 29.875 | 30 | 29.875 | 29.875 | 29.812 |
| 0.777778 | 29.875 | 29.938 | 30 | 29.875 | 29.938 | 29.812 |
| 0.784722 | 29.938 | 29.875 | 29.938 | 29.875 | 29.875 | 29.812 |
| 0.791667 | 29.875 | 29.938 | 30 | 29.875 | 29.938 | 29.812 |
| 0.798611 | 29.875 | 29.938 | 30 | 29.875 | 29.938 | 29.812 |
| 0.805556 | 29.875 | 29.938 | 30 | 29.938 | 30 | 29.875 |
| 0.8125 | 29.938 | 29.938 | 30.062 | 29.938 | 30 | 29.875 |
| 0.819444 | 29.938 | 29.938 | 30.062 | 29.938 | 29.938 | 29.812 |
| 0.826389 | 29.938 | 29.938 | 30 | 29.875 | 29.938 | 29.812 |
| 0.833333 | 29.938 | 29.938 | 30 | 29.938 | 29.938 | 29.875 |
| 0.840278 | 29.938 | 29.938 | 30 | 29.938 | 29.938 | 29.875 |
| 0.847222 | 29.938 | 29.875 | 30 | 29.938 | 29.938 | 29.75 |
| 0.854167 | 29.875 | 29.938 | 30 | 29.875 | 29.938 | 29.75 |
| 0.861111 | 29.875 | 29.875 | 29.938 | 29.812 | 29.938 | 29.75 |
| 0.868056 | 29.812 | 29.875 | 29.938 | 29.812 | 29.875 | 29.75 |
| 0.875 | 29.812 | 29.875 | 29.938 | 29.812 | 29.875 | 29.688 |
| 0.881944 | 29.812 | 29.875 | 29.938 | 29.812 | 29.875 | 29.688 |
| 0.888889 | 29.812 | 29.812 | 29.875 | 29.812 | 29.812 | 29.625 |
| 0.895833 | 29.75 | 29.75 | 29.875 | 29.75 | 29.812 | 29.625 |
| 0.902778 | 29.75 | 29.812 | 29.875 | 29.75 | 29.812 | 29.625 |
| 0.909722 | 29.75 | 29.688 | 29.875 | 29.688 | 29.812 | 29.562 |
| 0.916667 | 29.688 | 29.688 | 29.812 | 29.625 | 29.688 | 29.562 |
| 0.923611 | 29.688 | 29.625 | 29.75 | 29.625 | 29.688 | 29.5 |
| 0.930556 | 29.625 | 29.625 | 29.75 | 29.625 | 29.688 | 29.5 |
| 0.9375 | 29.625 | 29.562 | 29.75 | 29.625 | 29.625 | 29.5 |
| 0.944444 | 29.562 | 29.625 | 29.688 | 29.625 | 29.625 | 29.438 |
| 0.951389 | 29.5 | 29.625 | 29.75 | 29.562 | 29.625 | 29.438 |
| 0.958333 | 29.5 | 29.562 | 29.688 | 29.562 | 29.625 | 29.375 |
| 0.965278 | 29.562 | 29.562 | 29.688 | 29.562 | 29.562 | 29.438 |
| 0.972222 | 29.5 | 29.562 | 29.688 | 29.5 | 29.625 | 29.438 |
| 0.979167 | 29.5 | 29.562 | 29.688 | 29.562 | 29.625 | 29.375 |
| 0.986111 | 29.5 | 29.562 | 29.625 | 29.5 | 29.562 | 29.312 |
| 0.993056 | 29.5 | 29.5 | 29.625 | 29.5 | 29.5 | 29.312 |
| 1 | 29.5 | 29.5 | 29.562 | 29.5 | 29.562 | 29.25 |
| 1.00694 | 29.438 | 29.438 | 29.562 | 29.5 | 29.5 | 29.312 |
| 1.01389 | 29.438 | 29.438 | 29.562 | 29.438 | 29.5 | 29.25 |
| 1.02083 | 29.375 | 29.438 | 29.562 | 29.375 | 29.5 | 29.25 |
| 1.02778 | 29.438 | 29.375 | 29.562 | 29.438 | 29.438 | 29.188 |
| 1.03472 | 29.375 | 29.375 | 29.562 | 29.375 | 29.5 | 29.188 |
| 1.04167 | 29.375 | 29.438 | 29.562 | 29.375 | 29.438 | 29.188 |
| 1.04861 | 29.375 | 29.375 | 29.5 | 29.438 | 29.438 | 29.188 |
| 1.05556 | 29.312 | 29.438 | 29.5 | 29.375 | 29.438 | 29.188 |
| 1.0625 | 29.312 | 29.375 | 29.5 | 29.375 | 29.438 | 29.188 |
| 1.06944 | 29.438 | 29.438 | 29.562 | 29.438 | 29.5 | 29.312 |
| 1.07639 | 29.5 | 29.5 | 29.625 | 29.5 | 29.5 | 29.375 |
| 1.08333 | 29.5 | 29.562 | 29.688 | 29.625 | 29.562 | 29.375 |
| 1.09028 | 29.625 | 29.625 | 29.75 | 29.625 | 29.562 | 29.312 |
| 1.09722 | 29.625 | 29.688 | 29.75 | 29.562 | 29.562 | 29.375 |
| 1.10417 | 29.688 | 29.625 | 29.812 | 29.625 | 29.562 | 29.375 |
| 1.11111 | 29.688 | 29.688 | 29.812 | 29.688 | 29.688 | 29.375 |
| 1.11806 | 29.688 | 29.688 | 29.812 | 29.688 | 29.688 | 29.438 |
| 1.125 | 29.688 | 29.75 | 29.875 | 29.688 | 29.688 | 29.438 |
| 1.13194 | 29.75 | 29.75 | 29.875 | 29.688 | 29.688 | 29.438 |
| 1.13889 | 29.75 | 29.75 | 29.875 | 29.688 | 29.75 | 29.375 |
| 1.14583 | 29.688 | 29.75 | 29.875 | 29.75 | 29.75 | 29.438 |
| 1.15278 | 29.688 | 29.75 | 29.875 | 29.75 | 29.75 | 29.438 |
| 1.15972 | 29.75 | 29.75 | 29.875 | 29.688 | 29.75 | 29.375 |
| 1.16667 | 29.688 | 29.688 | 29.875 | 29.688 | 29.75 | 29.375 |
| 1.17361 | 29.688 | 29.688 | 29.875 | 29.688 | 29.75 | 29.312 |
| 1.18056 | 29.688 | 29.688 | 29.812 | 29.688 | 29.688 | 29.375 |
| 1.1875 | 29.688 | 29.75 | 29.875 | 29.688 | 29.75 | 29.375 |
| 1.19444 | 29.75 | 29.75 | 29.938 | 29.75 | 29.75 | 29.438 |
| 1.20139 | 29.75 | 29.75 | 29.938 | 29.812 | 29.75 | 29.438 |
| 1.20833 | 29.812 | 29.812 | 29.938 | 29.812 | 29.812 | 29.375 |
| 1.21528 | 29.75 | 29.812 | 29.938 | 29.812 | 29.812 | 29.438 |
| 1.22222 | 29.75 | 29.75 | 29.938 | 29.75 | 29.812 | 29.375 |
| 1.22917 | 29.75 | 29.75 | 29.875 | 29.75 | 29.75 | 29.375 |
| 1.23611 | 29.75 | 29.75 | 29.938 | 29.75 | 29.75 | 29.312 |
| 1.24306 | 29.688 | 29.75 | 29.938 | 29.75 | 29.75 | 29.312 |
| 1.25 | 29.688 | 29.75 | 29.938 | 29.75 | 29.75 | 29.312 |
| 1.25694 | 29.688 | 29.75 | 29.875 | 29.75 | 29.75 | 29.312 |
| 1.26389 | 29.688 | 29.688 | 29.938 | 29.812 | 29.75 | 29.375 |
| 1.27083 | 29.75 | 29.75 | 29.875 | 29.75 | 29.75 | 29.375 |
| 1.27778 | 29.75 | 29.812 | 29.875 | 29.75 | 29.75 | 29.312 |
| 1.28472 | 29.75 | 29.75 | 29.938 | 29.812 | 29.75 | 29.375 |
| 1.29167 | 29.812 | 29.75 | 30 | 29.812 | 29.812 | 29.375 |
| 1.29861 | 29.812 | 29.812 | 30 | 29.812 | 29.75 | 29.375 |
| 1.30556 | 29.875 | 29.812 | 30 | 29.875 | 29.812 | 29.438 |
| 1.3125 | 29.875 | 29.875 | 30 | 29.938 | 29.875 | 29.438 |
| 1.31944 | 29.875 | 29.938 | 30.062 | 29.938 | 29.875 | 29.438 |
| 1.32639 | 29.875 | 29.938 | 30.062 | 29.938 | 29.875 | 29.5 |
| 1.33333 | 29.938 | 29.938 | 30.125 | 30 | 29.938 | 29.438 |
| 1.34028 | 29.875 | 29.938 | 30.125 | 29.938 | 29.938 | 29.5 |
| 1.34722 | 29.938 | 30 | 30.188 | 29.938 | 29.938 | 29.5 |
| 1.35417 | 29.938 | 30 | 30.125 | 30 | 29.938 | 29.5 |
| 1.36111 | 30 | 30 | 30.188 | 30 | 30 | 29.5 |
| 1.36806 | 30 | 30.062 | 30.188 | 30.062 | 30 | 29.5 |
| 1.375 | 30 | 30 | 30.188 | 30 | 30 | 29.562 |
| 1.38194 | 30 | 30.062 | 30.188 | 30.062 | 30.062 | 29.562 |
| 1.38889 | 30.062 | 30.062 | 30.25 | 30.125 | 30 | 29.562 |
| 1.39583 | 30.062 | 30.062 | 30.25 | 30.125 | 30.062 | 29.562 |
| 1.40278 | 30.062 | 30.125 | 30.25 | 30.062 | 30.062 | 29.625 |
| 1.40972 | 30.125 | 30.125 | 30.312 | 30.188 | 30.125 | 29.625 |
| 1.41667 | 30.188 | 30.188 | 30.312 | 30.188 | 30.125 | 29.688 |
| 1.42361 | 30.188 | 30.188 | 30.375 | 30.188 | 30.188 | 29.688 |
| 1.43056 | 30.25 | 30.188 | 30.375 | 30.25 | 30.188 | 29.75 |
| 1.4375 | 30.25 | 30.25 | 30.375 | 30.25 | 30.25 | 29.688 |
| 1.44444 | 30.312 | 30.312 | 30.438 | 30.312 | 30.312 | 29.75 |
| 1.45139 | 30.312 | 30.375 | 30.5 | 30.312 | 30.312 | 29.812 |
| 1.45833 | 30.375 | 30.375 | 30.562 | 30.375 | 30.312 | 29.812 |
| 1.46528 | 30.375 | 30.375 | 30.562 | 30.375 | 30.438 | 29.875 |
| 1.47222 | 30.438 | 30.438 | 30.562 | 30.438 | 30.375 | 29.875 |
| 1.47917 | 30.438 | 30.438 | 30.562 | 30.438 | 30.438 | 29.875 |
| 1.48611 | 30.438 | 30.5 | 30.625 | 30.438 | 30.438 | 29.875 |
| 1.49306 | 30.438 | 30.438 | 30.625 | 30.438 | 30.438 | 29.875 |
| 1.5 | 30.5 | 30.5 | 30.688 | 30.5 | 30.5 | 29.938 |
| 1.50694 | 30.562 | 30.5 | 30.688 | 30.5 | 30.5 | 29.938 |
| 1.51389 | 30.562 | 30.562 | 30.75 | 30.562 | 30.5 | 30 |
| 1.52083 | 30.562 | 30.562 | 30.75 | 30.5 | 30.562 | 30 |
| 1.52778 | 30.625 | 30.625 | 30.75 | 30.562 | 30.625 | 30 |
| 1.53472 | 30.625 | 30.562 | 30.812 | 30.625 | 30.625 | 30 |
| 1.54167 | 30.625 | 30.625 | 30.812 | 30.625 | 30.625 | 30 |
| 1.54861 | 30.625 | 30.688 | 30.812 | 30.625 | 30.688 | 30.062 |
| 1.55556 | 30.688 | 30.688 | 30.812 | 30.688 | 30.688 | 30.062 |
| 1.5625 | 30.75 | 30.688 | 30.875 | 30.688 | 30.688 | 30.062 |
| 1.56944 | 30.688 | 30.75 | 30.875 | 30.688 | 30.688 | 30.125 |
| 1.57639 | 30.688 | 30.75 | 30.875 | 30.75 | 30.75 | 30.188 |
| 1.58333 | 30.75 | 30.75 | 30.938 | 30.75 | 30.75 | 30.125 |
| 1.59028 | 30.812 | 30.75 | 30.938 | 30.75 | 30.812 | 30.188 |
| 1.59722 | 30.812 | 30.812 | 31 | 30.812 | 30.812 | 30.188 |
| 1.60417 | 30.875 | 30.875 | 31 | 30.812 | 30.812 | 30.188 |

B

Temperature (°C)

| Fermentation time (days) | pH 6.5 | pH6 | pH5 | pH4 | pH3.5 |
| --- | --- | --- | --- | --- | --- |
| 0.173611 | 29.312 | 29.375 | 29.375 | 29.375 | 29 |
| 0.180556 | 29.375 | 29.562 | 29.438 | 29.438 | 29.062 |
| 0.1875 | 29.312 | 29.625 | 29.438 | 29.562 | 29.188 |
| 0.194444 | 29.375 | 29.688 | 29.5 | 29.625 | 29.25 |
| 0.201389 | 29.438 | 29.688 | 29.562 | 29.688 | 29.25 |
| 0.208333 | 29.438 | 29.75 | 29.562 | 29.75 | 29.312 |
| 0.215278 | 29.438 | 29.75 | 29.625 | 29.688 | 29.375 |
| 0.222222 | 29.438 | 29.688 | 29.625 | 29.688 | 29.375 |
| 0.229167 | 29.375 | 29.625 | 29.625 | 29.625 | 29.312 |
| 0.236111 | 29.438 | 29.562 | 29.562 | 29.562 | 29.312 |
| 0.243056 | 29.5 | 29.625 | 29.562 | 29.625 | 29.375 |
| 0.25 | 29.5 | 29.625 | 29.562 | 29.625 | 29.375 |
| 0.256944 | 29.5 | 29.625 | 29.625 | 29.625 | 29.375 |
| 0.263889 | 29.438 | 29.562 | 29.562 | 29.562 | 29.375 |
| 0.270833 | 29.438 | 29.5 | 29.562 | 29.562 | 29.375 |
| 0.277778 | 29.438 | 29.438 | 29.5 | 29.5 | 29.375 |
| 0.284722 | 29.5 | 29.562 | 29.562 | 29.625 | 29.375 |
| 0.291667 | 29.562 | 29.562 | 29.688 | 29.625 | 29.438 |
| 0.298611 | 29.562 | 29.625 | 29.688 | 29.625 | 29.438 |
| 0.305556 | 29.5 | 29.562 | 29.625 | 29.625 | 29.438 |
| 0.3125 | 29.562 | 29.625 | 29.625 | 29.625 | 29.5 |
| 0.319444 | 29.562 | 29.562 | 29.625 | 29.625 | 29.438 |
| 0.326389 | 29.562 | 29.562 | 29.625 | 29.625 | 29.438 |
| 0.333333 | 29.5 | 29.5 | 29.625 | 29.625 | 29.5 |
| 0.340278 | 29.5 | 29.562 | 29.625 | 29.625 | 29.5 |
| 0.347222 | 29.5 | 29.562 | 29.625 | 29.562 | 29.5 |
| 0.354167 | 29.5 | 29.5 | 29.625 | 29.562 | 29.375 |
| 0.361111 | 29.562 | 29.5 | 29.625 | 29.562 | 29.438 |
| 0.368056 | 29.562 | 29.5 | 29.625 | 29.562 | 29.438 |
| 0.375 | 29.5 | 29.5 | 29.562 | 29.562 | 29.438 |
| 0.381944 | 29.5 | 29.5 | 29.562 | 29.5 | 29.438 |
| 0.388889 | 29.5 | 29.5 | 29.562 | 29.562 | 29.438 |
| 0.395833 | 29.562 | 29.5 | 29.625 | 29.562 | 29.438 |
| 0.402778 | 29.5 | 29.562 | 29.562 | 29.562 | 29.438 |
| 0.409722 | 29.5 | 29.5 | 29.625 | 29.562 | 29.5 |
| 0.416667 | 29.5 | 29.5 | 29.562 | 29.562 | 29.5 |
| 0.423611 | 29.562 | 29.5 | 29.625 | 29.562 | 29.438 |
| 0.430556 | 29.5 | 29.5 | 29.625 | 29.562 | 29.438 |
| 0.4375 | 29.5 | 29.562 | 29.625 | 29.562 | 29.5 |
| 0.444444 | 29.562 | 29.5 | 29.625 | 29.562 | 29.5 |
| 0.451389 | 29.562 | 29.562 | 29.688 | 29.625 | 29.5 |
| 0.458333 | 29.562 | 29.562 | 29.625 | 29.625 | 29.562 |
| 0.465278 | 29.562 | 29.625 | 29.688 | 29.625 | 29.5 |
| 0.472222 | 29.562 | 29.562 | 29.688 | 29.625 | 29.5 |
| 0.479167 | 29.625 | 29.562 | 29.688 | 29.688 | 29.5 |
| 0.486111 | 29.625 | 29.625 | 29.688 | 29.688 | 29.562 |
| 0.493056 | 29.625 | 29.688 | 29.688 | 29.625 | 29.562 |
| 0.5 | 29.688 | 29.625 | 29.688 | 29.688 | 29.562 |
| 0.506944 | 29.625 | 29.625 | 29.75 | 29.688 | 29.562 |
| 0.513889 | 29.688 | 29.625 | 29.75 | 29.688 | 29.625 |
| 0.520833 | 29.688 | 29.688 | 29.75 | 29.75 | 29.562 |
| 0.527778 | 29.75 | 29.688 | 29.812 | 29.75 | 29.625 |
| 0.534722 | 29.688 | 29.688 | 29.75 | 29.75 | 29.688 |
| 0.541667 | 29.75 | 29.688 | 29.75 | 29.75 | 29.688 |
| 0.548611 | 29.75 | 29.688 | 29.812 | 29.75 | 29.625 |
| 0.555556 | 29.812 | 29.75 | 29.812 | 29.75 | 29.688 |
| 0.5625 | 29.75 | 29.688 | 29.812 | 29.75 | 29.625 |
| 0.569444 | 29.75 | 29.688 | 29.875 | 29.812 | 29.688 |
| 0.576389 | 29.812 | 29.75 | 29.875 | 29.75 | 29.688 |
| 0.583333 | 29.812 | 29.75 | 29.875 | 29.812 | 29.688 |
| 0.590278 | 29.812 | 29.75 | 29.875 | 29.812 | 29.688 |
| 0.597222 | 29.75 | 29.812 | 29.875 | 29.812 | 29.688 |
| 0.604167 | 29.75 | 29.75 | 29.875 | 29.812 | 29.688 |
| 0.611111 | 29.75 | 29.812 | 29.875 | 29.875 | 29.75 |
| 0.618056 | 29.812 | 29.812 | 29.875 | 29.812 | 29.75 |
| 0.625 | 29.812 | 29.75 | 29.875 | 29.875 | 29.75 |
| 0.631944 | 29.812 | 29.812 | 29.938 | 29.875 | 29.75 |
| 0.638889 | 29.875 | 29.812 | 29.938 | 29.875 | 29.75 |
| 0.645833 | 29.812 | 29.812 | 29.938 | 29.875 | 29.75 |
| 0.652778 | 29.875 | 29.875 | 29.938 | 29.875 | 29.812 |
| 0.659722 | 29.875 | 29.812 | 29.938 | 29.875 | 29.75 |
| 0.666667 | 29.875 | 29.812 | 29.938 | 29.938 | 29.75 |
| 0.673611 | 29.875 | 29.812 | 29.938 | 29.938 | 29.75 |
| 0.680556 | 29.875 | 29.812 | 30 | 29.875 | 29.812 |
| 0.6875 | 29.938 | 29.812 | 29.938 | 29.938 | 29.812 |
| 0.694444 | 29.875 | 29.875 | 30 | 29.938 | 29.812 |
| 0.701389 | 29.875 | 29.875 | 30 | 29.938 | 29.875 |
| 0.708333 | 29.938 | 29.875 | 30 | 29.938 | 29.812 |
| 0.715278 | 30 | 29.875 | 30 | 30 | 29.812 |
| 0.722222 | 30 | 29.875 | 30 | 29.938 | 29.812 |
| 0.729167 | 30 | 29.938 | 30 | 30 | 29.875 |
| 0.736111 | 30 | 29.875 | 30.062 | 30 | 29.812 |
| 0.743056 | 30 | 29.938 | 30 | 30 | 29.875 |
| 0.75 | 30 | 29.938 | 30.062 | 30 | 29.875 |
| 0.756944 | 30.062 | 29.938 | 30.125 | 30.125 | 29.875 |
| 0.763889 | 30.062 | 30 | 30.125 | 30 | 29.938 |
| 0.770833 | 30.062 | 30 | 30.125 | 30.062 | 29.938 |
| 0.777778 | 30.125 | 30 | 30.125 | 30.125 | 29.938 |
| 0.784722 | 30.125 | 30.062 | 30.188 | 30.125 | 29.938 |
| 0.791667 | 30.125 | 30.062 | 30.188 | 30.125 | 30 |
| 0.798611 | 30.188 | 30.062 | 30.188 | 30.188 | 30 |
| 0.805556 | 30.188 | 30.125 | 30.188 | 30.188 | 30 |
| 0.8125 | 30.188 | 30.125 | 30.25 | 30.188 | 30 |
| 0.819444 | 30.25 | 30.125 | 30.25 | 30.188 | 30 |
| 0.826389 | 30.25 | 30.125 | 30.25 | 30.188 | 30.062 |
| 0.833333 | 30.25 | 30.188 | 30.312 | 30.25 | 30.062 |
| 0.840278 | 30.312 | 30.188 | 30.375 | 30.25 | 30.062 |
| 0.847222 | 30.312 | 30.125 | 30.375 | 30.312 | 30.062 |
| 0.854167 | 30.375 | 30.188 | 30.312 | 30.312 | 30.062 |
| 0.861111 | 30.375 | 30.188 | 30.312 | 30.312 | 30.125 |
| 0.868056 | 30.375 | 30.25 | 30.312 | 30.312 | 30.125 |
| 0.875 | 30.375 | 30.25 | 30.375 | 30.312 | 30.188 |
| 0.881944 | 30.375 | 30.25 | 30.438 | 30.312 | 30.125 |
| 0.888889 | 30.375 | 30.188 | 30.375 | 30.312 | 30.062 |
| 0.895833 | 30.312 | 30.125 | 30.312 | 30.312 | 30.062 |
| 0.902778 | 30.25 | 30.125 | 30.312 | 30.188 | 30 |
| 0.909722 | 30.188 | 30.125 | 30.188 | 30.188 | 29.938 |
| 0.916667 | 30.125 | 30.062 | 30.188 | 30.188 | 29.938 |
| 0.923611 | 30.062 | 30 | 30.188 | 30.125 | 29.875 |
| 0.930556 | 30.125 | 29.938 | 30.125 | 30.062 | 29.812 |
| 0.9375 | 30 | 29.875 | 30.062 | 30 | 29.812 |
| 0.944444 | 30 | 29.875 | 30 | 30 | 29.75 |
| 0.951389 | 30 | 29.812 | 30 | 29.938 | 29.688 |
| 0.958333 | 30.125 | 29.938 | 30 | 30 | 29.75 |
| 0.965278 | 30.125 | 29.938 | 30.125 | 30.062 | 29.75 |
| 0.972222 | 30.125 | 30 | 30.062 | 30.125 | 29.75 |
| 0.979167 | 30.188 | 30.062 | 30.188 | 30.125 | 29.75 |
| 0.986111 | 30.188 | 30.062 | 30.188 | 30.062 | 29.75 |
| 0.993056 | 30.25 | 30 | 30.125 | 30.125 | 29.688 |
| 1 | 30.25 | 30.062 | 30.125 | 30.188 | 29.75 |
| 1.00694 | 30.25 | 30.125 | 30.188 | 30.188 | 29.812 |
| 1.01389 | 30.312 | 30.125 | 30.188 | 30.188 | 29.75 |
| 1.02083 | 30.25 | 30.125 | 30.25 | 30.188 | 29.812 |
| 1.02778 | 30.25 | 30.062 | 30.188 | 30.188 | 29.75 |
| 1.03472 | 30.188 | 30.125 | 30.188 | 30.188 | 29.75 |
| 1.04167 | 30.188 | 30.062 | 30.188 | 30.125 | 29.75 |
| 1.04861 | 30.188 | 30 | 30.125 | 30.125 | 29.688 |
| 1.05556 | 30.125 | 29.938 | 30.125 | 30.062 | 29.688 |
| 1.0625 | 30.125 | 29.938 | 30.125 | 30.062 | 29.688 |
| 1.06944 | 30.188 | 30 | 30.062 | 30 | 29.625 |
| 1.07639 | 30.125 | 29.938 | 30.062 | 30 | 29.625 |
| 1.08333 | 30.312 | 30.062 | 30.125 | 30.062 | 29.625 |
| 1.09028 | 30.375 | 30.125 | 30.188 | 30.125 | 29.75 |
| 1.09722 | 30.438 | 30.188 | 30.25 | 30.25 | 29.75 |
| 1.10417 | 30.438 | 30.25 | 30.312 | 30.25 | 29.75 |
| 1.11111 | 30.438 | 30.188 | 30.312 | 30.25 | 29.812 |
| 1.11806 | 30.438 | 30.188 | 30.25 | 30.25 | 29.75 |
| 1.125 | 30.438 | 30.188 | 30.312 | 30.25 | 29.75 |
| 1.13194 | 30.5 | 30.25 | 30.312 | 30.25 | 29.75 |
| 1.13889 | 30.562 | 30.312 | 30.375 | 30.438 | 29.875 |
| 1.14583 | 30.625 | 30.375 | 30.438 | 30.438 | 29.875 |
| 1.15278 | 30.625 | 30.375 | 30.438 | 30.438 | 29.938 |
| 1.15972 | 30.625 | 30.375 | 30.5 | 30.5 | 29.938 |
| 1.16667 | 30.625 | 30.375 | 30.5 | 30.5 | 29.938 |
| 1.17361 | 30.625 | 30.438 | 30.5 | 30.438 | 29.938 |
| 1.18056 | 30.688 | 30.438 | 30.5 | 30.5 | 29.938 |
| 1.1875 | 30.625 | 30.438 | 30.562 | 30.5 | 29.938 |
| 1.19444 | 30.562 | 30.438 | 30.5 | 30.438 | 29.875 |
| 1.20139 | 30.625 | 30.375 | 30.5 | 30.438 | 29.875 |
| 1.20833 | 30.75 | 30.438 | 30.5 | 30.5 | 29.938 |
| 1.21528 | 30.75 | 30.5 | 30.562 | 30.5 | 30 |
| 1.22222 | 30.75 | 30.5 | 30.625 | 30.562 | 29.938 |
| 1.22917 | 30.688 | 30.5 | 30.562 | 30.562 | 30 |
| 1.23611 | 30.688 | 30.438 | 30.625 | 30.5 | 29.938 |
| 1.24306 | 30.688 | 30.438 | 30.562 | 30.5 | 29.938 |
| 1.25 | 30.625 | 30.375 | 30.5 | 30.438 | 29.938 |
| 1.25694 | 30.562 | 30.375 | 30.5 | 30.438 | 29.875 |
| 1.26389 | 30.5 | 30.375 | 30.5 | 30.375 | 29.812 |
| 1.27083 | 30.5 | 30.312 | 30.438 | 30.375 | 29.812 |
| 1.27778 | 30.5 | 30.312 | 30.375 | 30.312 | 29.812 |
| 1.28472 | 30.562 | 30.312 | 30.438 | 30.375 | 29.812 |
| 1.29167 | 30.5 | 30.312 | 30.438 | 30.312 | 29.75 |
| 1.29861 | 30.5 | 30.312 | 30.438 | 30.312 | 29.75 |
| 1.30556 | 30.5 | 30.312 | 30.375 | 30.312 | 29.75 |
| 1.3125 | 30.5 | 30.312 | 30.438 | 30.312 | 29.75 |
| 1.31944 | 30.625 | 30.312 | 30.438 | 30.375 | 29.75 |
| 1.32639 | 30.75 | 30.5 | 30.5 | 30.438 | 29.875 |
| 1.33333 | 30.75 | 30.562 | 30.625 | 30.5 | 29.875 |
| 1.34028 | 30.812 | 30.625 | 30.688 | 30.562 | 29.938 |
| 1.34722 | 30.812 | 30.625 | 30.688 | 30.625 | 29.938 |
| 1.35417 | 30.875 | 30.625 | 30.75 | 30.625 | 29.938 |
| 1.36111 | 30.812 | 30.625 | 30.75 | 30.625 | 30 |
| 1.36806 | 30.812 | 30.625 | 30.75 | 30.688 | 30 |
| 1.375 | 30.875 | 30.688 | 30.75 | 30.688 | 30 |
| 1.38194 | 30.875 | 30.688 | 30.75 | 30.625 | 30 |
| 1.38889 | 30.875 | 30.625 | 30.75 | 30.625 | 30 |
| 1.39583 | 30.875 | 30.688 | 30.812 | 30.688 | 30.062 |
| 1.40278 | 30.875 | 30.75 | 30.812 | 30.688 | 30.062 |
| 1.40972 | 30.938 | 30.75 | 30.812 | 30.688 | 30.062 |
| 1.41667 | 30.938 | 30.75 | 30.812 | 30.75 | 30.062 |
| 1.42361 | 30.938 | 30.75 | 30.875 | 30.688 | 30.062 |
| 1.43056 | 30.875 | 30.75 | 30.875 | 30.75 | 30.125 |
| 1.4375 | 30.938 | 30.812 | 30.875 | 30.75 | 30.062 |
| 1.44444 | 30.938 | 30.75 | 30.875 | 30.75 | 30.125 |
| 1.45139 | 30.938 | 30.812 | 30.938 | 30.75 | 30.125 |
| 1.45833 | 30.938 | 30.812 | 30.938 | 30.75 | 30.188 |
| 1.46528 | 31 | 30.812 | 30.938 | 30.75 | 30.188 |
| 1.47222 | 31 | 30.812 | 30.938 | 30.812 | 30.188 |
| 1.47917 | 31 | 30.812 | 30.938 | 30.812 | 30.188 |
| 1.48611 | 31 | 30.875 | 31 | 30.812 | 30.188 |
| 1.49306 | 31 | 30.875 | 31 | 30.875 | 30.25 |
| 1.5 | 31 | 30.875 | 31 | 30.875 | 30.188 |
| 1.50694 | 31.062 | 30.938 | 31.062 | 30.812 | 30.25 |
| 1.51389 | 31.125 | 30.938 | 31.125 | 30.938 | 30.25 |
| 1.52083 | 31.125 | 30.938 | 31.125 | 30.938 | 30.25 |
| 1.52778 | 31.125 | 30.938 | 31.125 | 30.938 | 30.312 |
| 1.53472 | 31.125 | 31 | 31.188 | 30.938 | 30.312 |
| 1.54167 | 31.125 | 31 | 31.125 | 30.938 | 30.312 |
| 1.54861 | 31.188 | 31 | 31.188 | 31 | 30.312 |
| 1.55556 | 31.188 | 31.062 | 31.188 | 31 | 30.312 |

**Supplementary material 4 |** Ergosterol (A) and glucosamine (B) concentration (g/100 g DW) in pure mycelium of *S. rugosoannulata* (red), *S. commune* (orange), *P. cinnabarinus* (green), *V. volvacea* (turquoise), *P. pulmonarius* (blue) or *R. microsporus* (pink).

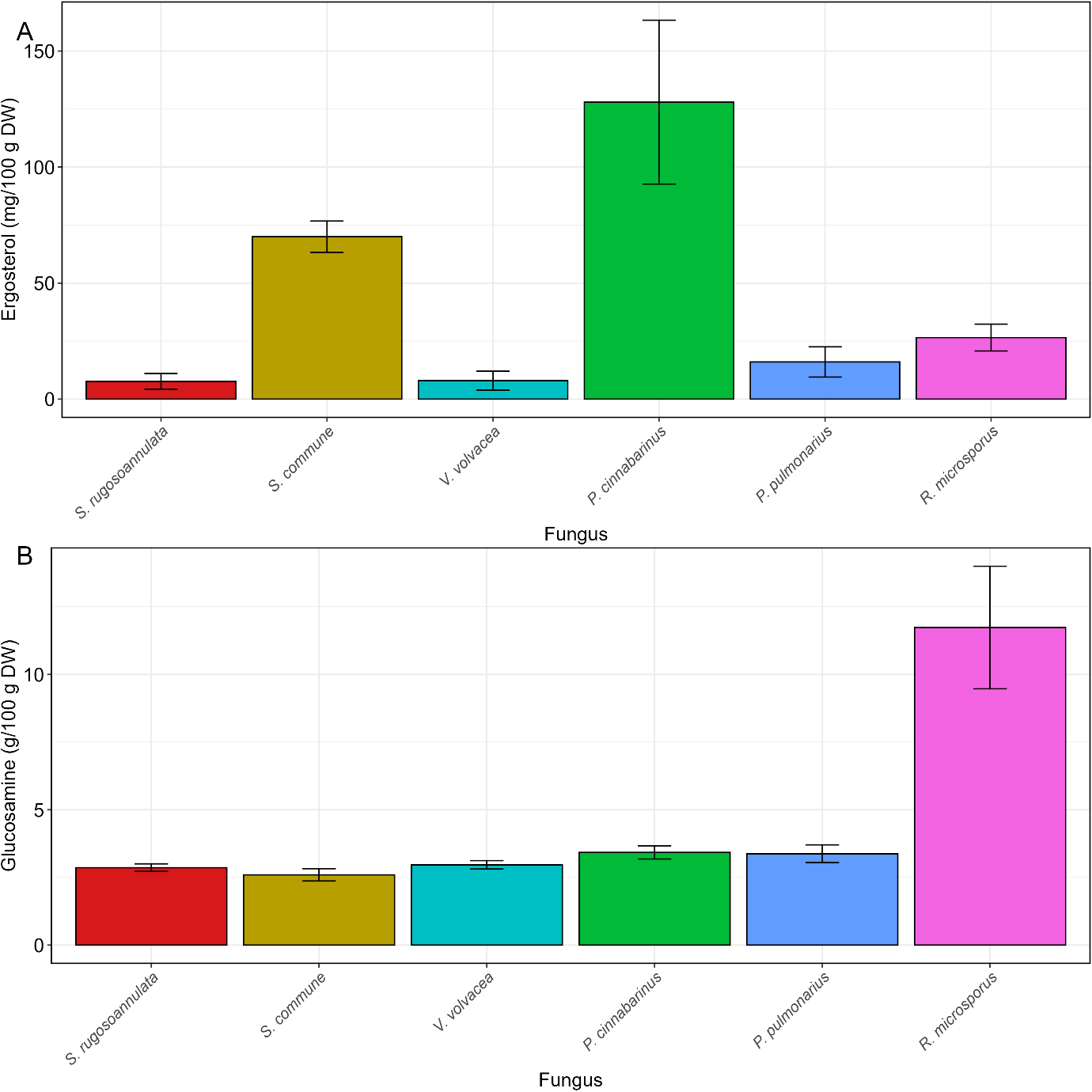

**Supplementary material 5 |** Correlation between fungal biomass based on glucosamine and ergosterol (g/100 g DW) in *R. microsporus* fermentations. R2 and significance in the top left corner.

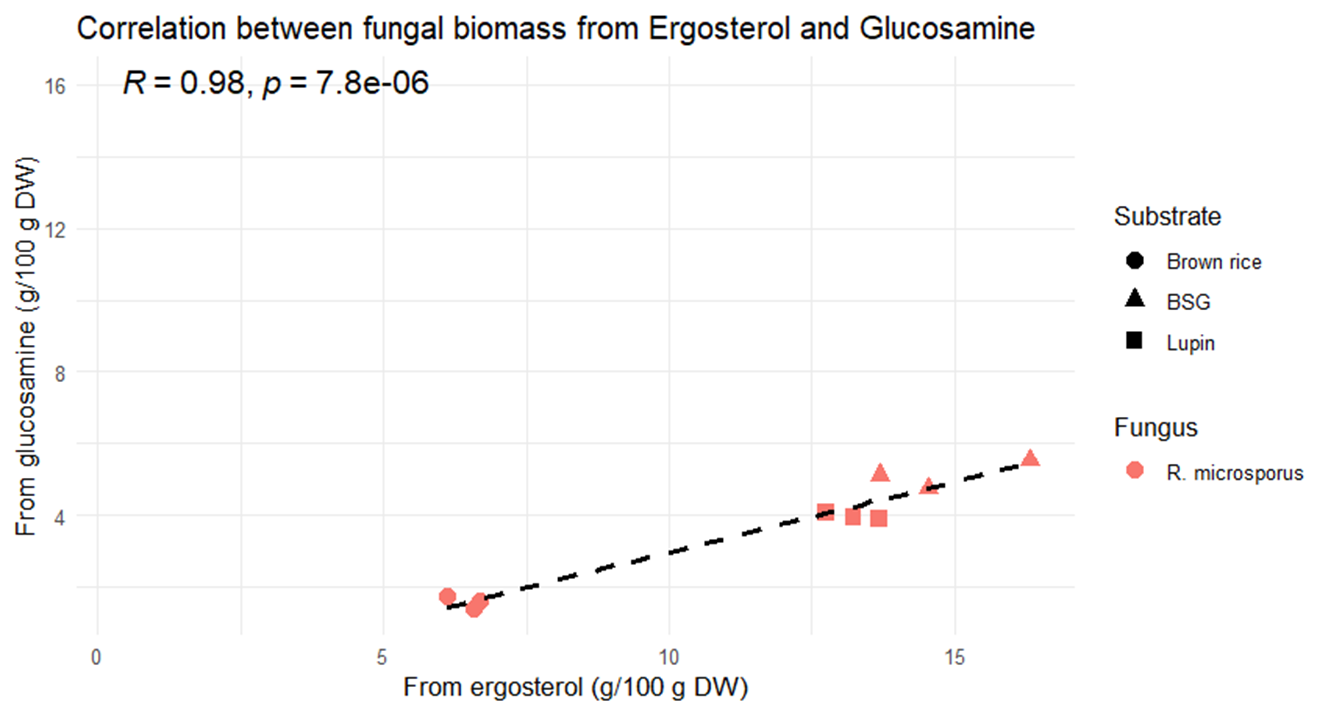
